# A multiscale analysis of DNA phase separation: From atomistic to mesoscale level

**DOI:** 10.1101/375626

**Authors:** Tiedong Sun, Alexander Mirzoev, Vishal Minhas, Nikolay Korolev, Alexander P. Lyubartsev, Lars Nordenskiöld

## Abstract

DNA condensation and phase separation is of utmost importance for DNA packing *in vivo* with important applications in medicine, biotechnology and polymer physics. The presence of hexagonally ordered DNA is observed in virus capsids, sperm heads and in dinoflagellates. Rigorous modelling of this process in all-atom MD simulations is presently difficult to achieve due to size and time scale limitations. We used a hierarchical approach for systematic multiscale coarse-grained (CG) simulations of DNA phase separation induced by the three-valent cobalt(III)-hexammine (CoHex^3+^). Solvent-mediated effective potentials for a CG model of DNA were extracted from all-atom MD simulations. Simulations of several hundred 100-bp-long CG DNA oligonucleotides in the presence of explicit CoHex^3+^ ions demonstrated aggregation to a liquid crystalline hexagonally ordered phase. Following further coarse-graining and extraction of effective potentials, we conducted modelling at mesoscale level. In agreement with electron microscopy observations, simulations of an 10.2-kbp-long DNA molecule showed phase separation to either a toroid or a fibre with distinct hexagonal DNA packing. The mechanism of toroid formation is analysed in detail. The approach used here is based only on the underlying all-atom force field and uses no adjustable parameters and may be generalized to modelling chromatin up to chromosome size.

## INTRODUCTION

The compaction of DNA is a problem of outstanding importance in biology with many important applications in polyelectrolyte theory, biotechnology, nanoscience (1–4). While a long (~100 Mbp) chromosomal DNA molecule in low salt solution would adopt a random coil conformation expanding over 100 μm, 46 such DNA molecules are packed inside the confined space of about 10 μm in the human cell nucleus. Similarly, in sperm heads, viruses and bacteria, DNA is extremely densely condensed (4–7). The packaging of the giant genomes of dinoflagellates is another example of compact ordered liquid-crystalline form of DNA (8,9). Notably, in the eukaryotic cell nucleus formation of heterochromatin has recently been proposed to proceed through liquid-liquid phase separation of condensed DNA domains (10,11). DNA condensation has also attracted great attention in gene delivery where the compaction is a key to optimizing DNA transfer (3,12).

For almost 50 years it has been known that *in vitro*, in the presence of highly charged cations like cobalt(III)-hexammine (CoHex^3+^), spermidine^3+^ and spermine^4+^, DNA in solution condenses into collapsed structures of varying morphologies such as toroids, rod like fibres, globules and liquid crystals (13–17). While liquid crystalline phases are observed for 150 bp or shorter DNA molecules (18–20), long DNA molecules (a few to several hundred kbp) exhibit highly regular toroidal structures with DNA arranged in hexagonal packing inside the toroids (6), which have an outer diameter of around 100 nm, depending on conditions (21,22). This spontaneous formation of DNA toroids in hexagonal arrangement is also observed *in vivo* in viruses and sperm chromatin and has fascinated scientists for a long time (5,6,23,24). This phenomenon has been vastly studied experimentally with a variety of techniques such as X-ray diffraction (25), Cryo-EM (22,24) and more recently with single molecule techniques (26). This has resulted in significant advances in our understanding of the phenomenon both at mechanistic and fundamental level. However, there are still many unsolved problems related to the condensation of DNA induced by multivalent cations resulting in the formation of the ordered DNA liquid crystalline phase, and the collapse of DNA into toroidal structures. There is a lack of rigorous theoretical modelling approaches that are able to predict, reproduce and analyse these phenomena from basic principles.

Although it may seem counterintuitive, the fundamental origin of multivalent ion induced attraction between like-charged DNA molecules leading to condensation is grounded in the electrostatic properties of the highly charged DNA polyelectrolyte. Based on computer simulations as well as on analytical theories, it has been established that the attraction is caused mainly by ion-ion correlations that result in a correlated fluctuation in the instantaneous positions of the condensed counterions on DNA, leading to a net attractive force between DNA molecules (reviewed in (2,27)). In the case of flexible multivalent cations like the polyamines or oligopeptides, the attraction is also generated by the “bridging” effect (28). The origin of multivalent ion induced attraction between aligned DNA-DNA molecules is therefore clear and well described by such polyelectrolyte models, and recently also in all-atom detail (29,30). However, the mesoscale level spontaneous transition of short DNA molecules to a hexagonal liquid crystalline phase and the formation of toroids from kbp-long DNA molecules, have not been rigorously described and analysed theoretically. These phenomena that are determined by a combination of electrostatic forces and by DNA mechanical properties, are highly important for understanding DNA phase separation in biology.

Computer modelling of DNA condensation to an ordered bundled phase or modelling of single DNA molecule collapse to the toroid structure, have with few exceptions been performed with a description of DNA as a chain of beads using parameterised harmonic bonds. The bending flexibility was commonly tuned to reproduce the DNA mechanical properties from persistence length data (31–34). Furthermore, the DNA-DNA attraction was generally modelled by empirical potentials. Only in a few of these works electrostatic effects were treated directly by including explicit balancing counterions, however, without added salt (31–33). Common to all these approaches is that they treat generic polymer molecules without explicit presence of added salt, using empirical adjustable parameters to describe the relevant potentials in the models. Hence, the connection to the atomistic DNA structure and chemical specificity is lost. The experimentally important phenomenon of the formation of hexagonally ordered liquid crystalline phase of short DNA molecules in the presence of multivalent counterions has to the best of our knowledge never been theoretically demonstrated.

In recent years advances in computer technology have progressed considerably and all-atom biomolecular MD simulations including molecular water can now be performed for very large systems such as a nucleosome core particle and large DNA assemblies (30,35). Yoo and Aksimentiev developed improved ion-phosphate interaction force field parameters and performed all-atom MD simulations of an array of 64 parallel duplex DNA. They demonstrated the correct hexagonal packing of DNA, which was absent in simulation with standard CHARMM or AMBER force field parameters (36). The same authors also investigated the physical mechanism of multivalent ion-mediated DNA condensation at atomistic level. The results supported a model of condensation driven by entropy gain due to release of monovalent ions and by bridging cations (30).

However, all-atom MD simulation of DNA liquid crystalline formation or kbp-size DNA toroid condensation is presently not computationally feasible and hence multiscale approaches linking atomistic and coarse-grained (CG) levels of description are necessary (37). Within a systematic bottom-up multiscale modelling scheme, the macromolecules are reduced to a CG description with effective sites representing groups of atoms and a number of such coarse-graining methods and models for DNA have been developed, each with respective strengths and weaknesses (38).

Some bottom-up approaches, have recently been used to model DNA (39–42) and various DNA properties (flexibility, topological effects, interaction with proteins, DNA aggregation (43–45); see also reviews (37,46)). Most of the reported studies did not treat both electrostatics and solvent effects rigorously, i.e. did not use explicit ion models, or did not study the mesoscopic scale of DNA condensation. An alternative to systematic bottom-up CG approach for modelling DNA at mesoscale level is represented by recent work by the de Pablo group using a top-down approach to fit the model parameters to experimental data (DNA thermal denaturation) (41,47). The DNA model combined with explicit ions was used in simulations of 4 kbp DNA packaging in the presence of multivalent ions inside a virus capsid (48).

The DNA condensation to a hexagonally ordered phase as well as DNA toroid formation, both induced by the presence of multivalent cations, are phenomena clearly intrinsic to the DNA molecule and inherent in its physico-chemical properties. We hypothesize that this behaviour can be predicted by a bottom-up approach that is based on state-of-the-art all-atom molecular dynamics (MD) simulations, which is followed by structure-based coarse-graining resulting in effective interaction potentials for the CG model without further adjustable parameters.

Here, we perform systematic multiscale structure-based CG simulations of DNA with explicit electrostatic interactions included, starting from all-atom description going up to mesoscale modelling of DNA. The CG model is simple enough yet captures the structure form of double helix DNA and rigorously treats the electrostatic interactions. The systematic coarse-graining follows the inverse Monte Carlo (IMC) approach to extract solvent mediated effective CG potentials for all interactions in the system (49,50) from structural properties of the underlying system. The model is validated in CG simulations of DNA persistence length as a function of monovalent salt demonstrating outstanding performance. We show DNA condensation induced by the three-valent CoHex^3+^ ion for short DNA duplexes resulting in a bundled phase with hexagonal ordering. Furthermore, adopting a second level “super coarse-grained” (SCG) DNA beads-on-a-string model we show that this approach remarkably predicts the hexagonally ordered liquid crystalline-like phase of short DNA and toroid formation in hexagonal arrangement for kbp-size long DNA, giving mechanistic insight on the DNA condensation process. In order to pave the ground for detailed analysis of the compaction of DNA at mesoscale level in chromosomes, it is necessary to develop similarly chemically informed bottom-up models for the amino acids modelling histone proteins. Such models should have the predictive power to be trusted in modelling biologically important phenomena at mesoscale level for experimentally unexplored scenarios. The present approach represents a first and successful step in this direction.

## METHODS

More details on computational methods and parameters used in this work are given in the Supplementary Methods section of the Supplementary Data (SD).

### Hierarchy of DNA models

The hierarchical multiscale approach used here implements systematic stepwise coarse-graining of DNA model from the atomistic to a beads-on-string level as illustrated in Figure 1. Coarse-graining is performed at two spatial scales resulting in three DNA models with different resolutions. Figure 1A illustrates the all-atom DNA model which includes explicit water molecules and ions. In the CG DNA model (Figure 1B), the DNA is modelled with five beads, representing a two-base pair unit of DNA (Figure 1D). There are four types of bonds and three types of angles in the bonded interaction terms (Figure 1D). We include explicit mobile ions and charges of DNA phosphate groups while the solvent is considered implicitly. As detailed below and in the Supplementary Data, for the CG model we have performed three different parameterisations at the CG level, based on three different all-atom systems.

**Figure 1.**
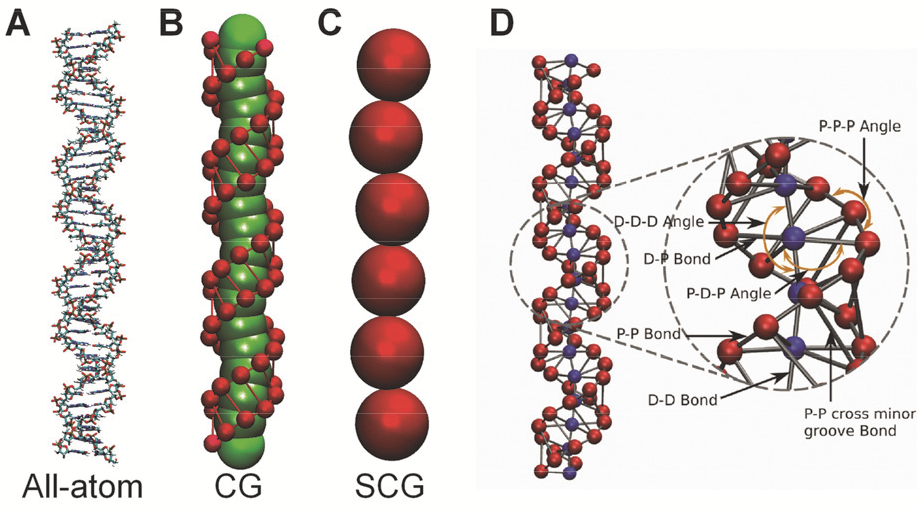
Hierarchy of DNA models. (**A**) The all-atom DNA model. (**B**) The first level of DNA coarse-graining with two bead types. (**C**) The SCG model is built upon the CG model with one bead type. (**D**) The detailed presentation of the CG DNA model in (**B**). Bond interactions, D-P, P-P (backbone), D-D and P-P (cross minor groove) are shown in grey. Angle interactions (P-D-P, P-P-P and D-D-D) are indicated by orange arrows.

For the CG model, this results in a large number of interactions parameterised, all of them calculated by the IMC procedure except for long-range electrostatics. The exact number of interactions depends on the composition of the CG system. In summary, the CG model includes the following type of interactions: i) bonded interactions between DNA beads (see Figure 1D), ii) angle interactions between DNA beads (Figure 1D), iii) intermolecular short-ranged non-bonded terms between DNA beads, iv) intermolecular short-ranged non-bonded terms between charged mobile ions as well as between ions and DNA D/P beads, v) long-range electrostatic interactions, which are the same in all systems and described by Coulombic interactions in a dielectric continuum described by the dielectric permittivity of water, vi) short-range intramolecular interactions within one DNA molecule (see details in the Supplementary Methods). Table S1 summarises all interaction types for the three different CG parameterisations performed (see details below and in the Supplementary Data).

The second level of coarse-graining results in a beads-on-string type of model, called the super CG DNA (SCG) model, which is shown in Figure 1C. Here, a single type of uncharged beads (called ”S”) represents three units of the CG DNA model, corresponding to six DNA base pairs. There is one bond and one angle potential as well as the presence of non-bonded intermolecular and intramolecular effective potentials between the S beads, which are parameterised by IMC. In the SCG model both solvent and electrostatic interactions are implicitly included into the effective potentials (see details below).

It may also be noted that the geometry and topology of the CG model preserves the DNA chirality. However, this model would also be applicable to a left-handed helix, but under the present conditions, the structure is constrained to the initial right-handed form due to the large conversion barrier. The DNA in the SCG model does not have chirality.

### All-atom Molecular Dynamics simulations

Three different all-atom simulation corresponding to set-up of the given all-atom system are set up. These correspond to the three parametrisations made within the IMC procedure (see details below).

First, we perform all-atom MD simulations of a single 40 bp DNA in the presence of 130 mM NaCl. This is used in IMC parametrisations for validation of experimental behaviour of persistence length as a function of ionic strength.

Second, simulations in the presence of CoHex^3+^ ions, which induces condensation of DNA to fibres are performed. Here, three independent microsecond-long all-atom MD simulations are performed with four 36 bp-long DNA molecules (the sequences are given in the SD) in presence of water and ions as illustrated in Figure 2. The system contains CoHex^3+^ ions corresponding to 1.5 times the charge of the DNA. Additionally the system contains added salt corresponding to 50 mM K^+^ and 35 mM Na^+^, with neutralizing amount of Cl^−^ ions. The CHARMM27 force field is used, supplemented with Car-Parrinello MD (CPMD) optimized CoHex^3+^ parameters obtained in our previous work (51).

**Figure 2.**
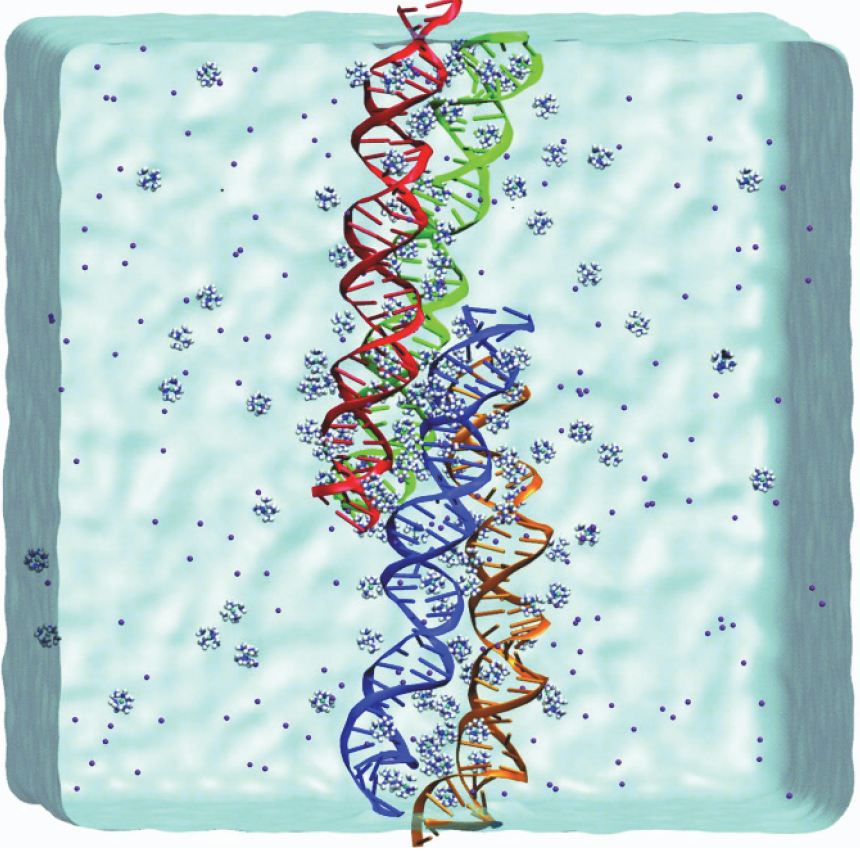
Bundling of DNA in all-atom MD simulations in the presence of CoHex^3+^. A snapshot shows formation of a bundle of the four 36 bp DNA double helices (coloured blue, red, green and orange) in the cubic box of 15 nm size. Ions (140 CoHex^3+^, 140 K^+^, 95 Na^+^, 375 Cl^−^) are shown as spheres and explicit water is displayed as blue surface.

Third, a similar setup as case two above, but with the CoHex^3+^ ions replaced by Mg^2+^ is made. This serves as a control simulation performed to demonstrate absence of DNA-DNA attraction and no DNA condensation in all-atom and in CG simulations in the presence of Mg^2+^ as observed in experiments.

The CHARMM27 force field is used in all simulations, supplemented with Car-Parrinello MD (CPMD) optimized CoHex^3+^ parameters obtained in our previous work (51).

### Inverse Monte Carlo procedure

In both the CG and SCG DNA models, all effective interaction potentials are obtained in a systematic and rigorous way using the IMC method (49,50). The potentials are constructed to reproduce selected average structural properties of the fine-grained system such as radial distributions functions (RDF) between non-bonded CG sites and bond length and angle distributions for bonded sites. For the CG DNA model, these are obtained by mapping the all-atom MD trajectory from the four-DNA system to a corresponding CG site representation. Thus, no empirical or adjustable parameters are used in the CG DNA model and it rests only on the parameters from all-atom CHARMM27 DNA as well as on the CPMD optimized parameters for CoHex^3+^. The IMC-derived potentials are tabulated and not prescribed by any specific functional form.

The topology of the CG DNA model (Figure 1B and 1D) is similar to our previous CG DNA model (52,53) but with all interaction potentials derived by the IMC method from the underlying atomistic simulations. The total number of bead types is eight: four DNA beads (D, P, as well as terminal DT, PT), one CoHex^3+^, one K^+^, one Na^+^ and one Cl^−^ bead. The complete set of interaction parameters consists of 36 non-bonded terms, 4 bonded terms and 3 angle terms (see details in Supplementary Data, Table S1). All effective interaction potentials are derived simultaneously, so that all correlations between different interaction terms are taken into account. It can be noted that the present DNA model is not parameterised for sequence specificity but such an extension is possible by treating each unique two-base pair step individually at the mapping stage (there being 10 unique steps).

The potentials for the SCG model are derived in the same way as for the CG model using the IMC method. Trajectories generated by MD simulation of the CG model are mapped to the SCG representation. Then the RDF between the “S” sites, as well as bond and angle distributions are calculated with the mapped SCG trajectories.

It should be noted that in addition to the short-range part of effective potential (which has cut-off range of 25 Å), all charged sites are interacting by a Coulombic potential (scaled by the dielectric constant). This means that beyond the range of 25 Å DNA, interactions are primarily electrostatic-driven. These long-range interactions are treated using Ewald summation. The dielectric constant is set to 78.0. A detailed discussion of this approach is given in (54).

Effective ionic potentials for the same interactions that are obtained from different parameterisations are generally very similar (see below). However, our methodology is structure-based, and it is therefore preferable to make separate parameterisations for systems that display considerably different structural features in experimental behaviour and conditions.

In summary, we have performed three different parameterisations at the CG level, based on three different all-atom systems. General information about these three levels of simulations and their computational efficiency (CPU usage time) is given in Table S2 in the SD. Table S1 lists all the interaction potentials that are calculated in the CG models for the three separate parameterisations that have been performed. All tabulated potentials are provided in the SD in an archive zip file.

The IMC calculation is carried out with the MagiC software v.3 (55) that is also used for bead-mapping, RDF calculation, analysis and export of the resulting potentials.

### Coarse-grained simulations

Following the extraction of effective CG potential with the IMC method, we use these potentials to perform CG MD simulations for a system comprising two hundred DNA molecules and explicit ions to simulate DNA aggregation in the presence of CoHex^3+^. This simulation is used for further coarse-graining to a “super-CG” (SCG) DNA model (Figure 1C) with another step of IMC. The simulations with the CG and SCG DNA models are conducted using the LAMMPS package (56) within the NVT ensemble. CG model simulations were carried out for 200 pieces of 100-bp DNA (50 CG-DNA units) in a 150×150×150 nm^3^ simulation box containing 13,200 CoHex^3+^ ions (corresponding to a concentration of 6.5 mM) in the presence of 10 mM K^+^ and 10 mM Na^+^ ions as well as neutralizing Cl^−^ ions. The derived effective potentials for the SCG model enable us to simulate DNA condensation and phase separation at mesoscale level. We perform simulations for two systems. The first comprises 400 pieces of 96 bp DNA (represented by a chain of 16 S-beads) and the second system is a single 10.2 kbp-long DNA (1700 S-beads). Within the SCG model, the electrostatic interactions are treated implicitly since they are effectively included into the SCG effective potentials.

### Persistence length calculation

To validate the CG DNA model we test its performance in prediction of the salt dependence of the DNA persistence length, *L_p_*. We perform all-atom MD simulations of a single 40 bp DNA in the presence of physiological salt (130 mM NaCl). Following that we make the mapping of this trajectory to the CG model and extract effective IMC potentials. To estimate the dependence of the persistence length on ionic concentration, a 500 bp long CG DNA is simulated in the range of NaCl salt concentrations from 0.1 mM to 100 mM. Our simulation results are compared with experimental data and with data from other simulation studies.

The persistence length is calculated according to the formula *L_p_* = −*L_c_*/(ln(<*cos α*>)). Here, *L_c_* is the contour length of the DNA fragment and <*cos α*> is the average of the cosine between two adjacent DNA segments. The average is taken over the length of the simulation and over the positions of the segments on DNA for the corresponding contour length. Then (ln(<*cos α*>) is plotted as a function of *L_c_*, and *L_p_* is estimated by determining the slope of the plot, as done in our previous study (53). We do not determine *L_p_* in atomistic simulations, but in coarse-grained simulations, where the length of DNA (500 bp) is longer than the Kuhn segment.

Physically, the persistence length is defined by the DNA local flexibility and by the long-range interactions which have mostly electrostatic and excluded volume nature. The DNA local flexibility is determined by the bonded interactions of the CG model, which are well parametrized from the atomistic simulations of the 40 bp fragment. The long-range interactions of the CG model consist of a short-range part (within cut-off distance) representing excluded volume and solvent-mediated interactions, which is parametrized from the atomistic simulations, and the electrostatic part which is not limited by distance but strictly treated in the CG simulations by the Ewald method.

## RESULTS AND DISCUSSION

### Validation of the CG DNA model and the approach

First we validate the approach and the CG DNA model (described above) against experimental persistence length data as a function of monovalent salt concentration. We perform an all-atom MD simulation of a single 40 bp DNA in the presence of physiological salt (130 mM NaCl). The all-atom trajectory is run for 2 μs and demonstrates stability of helical parameters and the elasticity of the DNA double helix (data not shown). We then extract effective short-range potentials (see Supplementary Methods for details). Figure S1 of the Supplementary Data (SD) shows all the effective potentials. Following that, we proceed to run several CG simulations of a single 500 bp-long DNA molecule in the presence of varying concentration of monovalent salt in the range 0.1 to 100 mM. Figure 3 compares the dependence of persistence length as a function of salt concentration with experimental data (57–63). We include experimental persistence length data from several sources as there is a large variation in these results depending on the method used and the procedure for analysing the original measurements. Additionally, we also include values predicted by other CG DNA models with explicit ions, obtained using bottom-up approaches based on underlying all-atom MD simulations of single DNA oligonucleotides (41,42,64).Considering the variation in experimental data, the present model demonstrates excellent agreement with experiments for three orders of magnitude variation in salt concentration. However, the results of our model seem to display somewhat lower *L_p_* values as compared to experiment, which implies that within this model, DNA may be slightly more flexible than real DNA. In general, the performance of our model is superior to other available explicit ion CG DNA models. This accurate prediction of the effect of electrostatic interactions on DNA flexibility lends confidence to the present approach. The result shows that the model well represents both DNA intrinsic flexibility and electrostatic interactions. A more detailed discussion of DNA flexibility and persistence length prediction from the CG model and how the DNA flexibility depends on choice of underlying force field will be presented elsewhere (Minhas *et al*., to be published).

**Figure 3.**
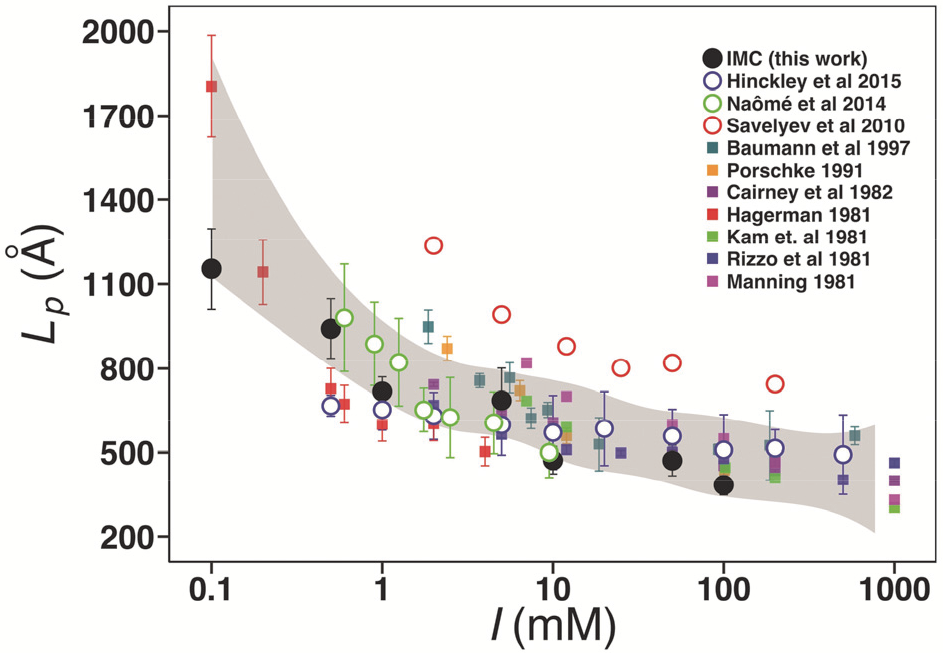
Dependence of DNA persistence length (*L_p_*) on NaCl concentration (*I*). Solid black circles are data of this work. Solid squares are experimental results: dark cyan (57), orange (58), dark magenta (59), red (60), green (61), blue (62), magenta (63). Hollow circles are results from other computer simulations with explicit ions: blue (41), green (64), red (42). Shaded area is the spline approximation of the experimental data with confidence level 0.99995.

### DNA-DNA attraction in all-atom MD simulations

We perform all-atom MD simulations for a system with four DNA molecules containing explicit ions (CoHex^3+^, Na^+^, K^+^ and Cl^−^) and water. A snapshot of the system is shown in Figure 2. Data from the three independent 1 μs-long simulations are subsequently used in the IMC procedure for extraction of effective solvent mediated potentials for the CG DNA model.

Similarly to the result of our previous work (51), the system shows DNA-DNA attraction and aggregation of DNA into fibre-like bundles induced by CoHex^3+^ (Figure 2). DNA fibres are formed across the periodic boundaries in all three independent simulations (see Supplementary Figure S2, AC). The snapshots show some variability in the character of DNA-DNA contacts over the periodic images. In order to obtain rigorous RDFs it is important to analyse an equilibrated ensemble. It is, however, clear that for an all-atom MD simulation of a system of molecules of this size with very strong electrostatic coupling, it cannot be claimed that the system is fully equilibrated. This is the reason why we averaged the RDF from 3 independent runs. We believe that typical configurations of several local minima sampled in independent simulations can capture the most characteristic configurations of DNA and ions, providing a realistic estimate of the intermolecular interactions. Furthermore, the electrostatics is the main driving force driving the large scale behaviour of polyelectrolytes such as DNA, while the specific short-range interactions determine details (65). It is therefore not unreasonable to suggest that these short-range specific interactions, which are determined by RDFs obtained from atomistic simulations can be realistically reproduced by sampling configurations within local energy minima. Since CG models generally present a smoother free energy surface than their fine-grained counterpart, more efficient sampling can be achieved in CG MD simulations. It should, however, be noted that the condensed configurations may not represent the lowest free energy state, being constrained by the local minimum.

After extracting the effective potentials for the CG model, longer CG MD simulations of the same system with four DNA molecules, are performed (see more details in the next section). A final snapshot from one CG-MD simulation of this four DNA system is shown in the Supplementary Figure S2D. In the CG MD simulations a straight fibre configuration is easily reached. This result is stable and does not vary in different simulations. This gives additional confidence for the trust in the effective potentials obtained from the all-atom MD simulation, in spite of the general problem of sampling of the complete configuration space in the all-atom simulations.

### Coarse-Grained DNA model with rigorous effective solvent mediated potentials

Following the all-atom MD simulations of the system with four DNA molecules in all-atom representation containing explicit ions (CoHex^3+^, Na^+^, K^+^ and Cl^−^) and water as described above, we proceed to extract the effective potentials for the CG model. The trajectories generated by the three atomistic MD simulations are mapped from the all-atom to the CG DNA model. We use these for calculation of RDFs and intramolecular (within DNA) distributions of bond lengths and angles between the CG sites, with averaging over all three independent trajectories. Examples of calculated RDFs are shown in Figure 4A-C. Selected distribution functions and effective potentials are plotted in Figure 4A-F. All RDFs and effective potentials for the CG DNA model in the presence of CoHex^3+^ can be found in Supplementary Figure S3. All charged sites interact by a Coulombic potential (scaled by the dielectric constant) and treated by the Ewald summation. DNA interacts by this potential beyond 25 Å, which is the cut-off used only for the short-range part of the total potential. It is only the short-ranged parameterised part of the potential that is displayed in Fig 4. Supplementary Figure S4 illustrates convergence in the IMC calculations.

**Figure 4.**
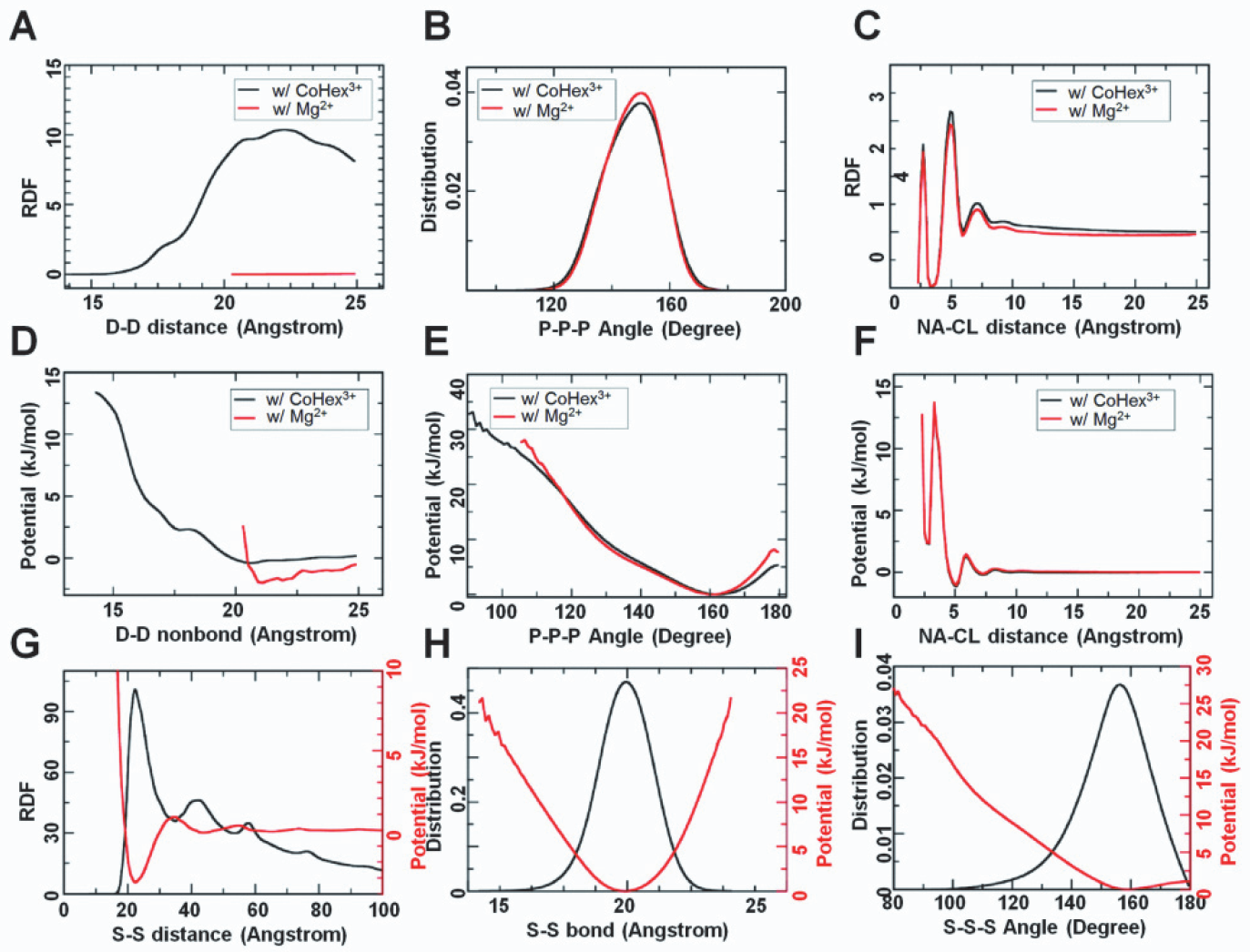
Illustration of RDFs and corresponding effective potentials obtained by the IMC method. Selected RDFs (top row), effective potentials of the CG model (middle row) and all RDFs and effective potentials of the SCG model (bottom row). In (**A-F**), three distributions and potentials are presented: D-D non-bonded pair, P-P-P angle along backbone and Na-Cl non-bonded pair, from the systems with CoHex^3+^ (black) and Mg^2+^ (red) ions. In (**G-I**), distributions and effective potentials of the SCG model are plotted together for the S-S non-bonded interaction (**G**), the S-S bond (**H**) and the S-S-S angle (**I**)

We additionally performed a control all-atom MD simulation without the presence of CoHex^3+^ under conditions that should not lead to DNA condensation. The CoHex^3+^ ions are replaced by the same number of Mg^2+^ ions (with the corresponding decrease in number of Cl^−^ counterions), resulting in the absence of DNA condensation, which is in agreement with the experimentally known fact that the divalent Mg^2+^ does not induce DNA condensation (1) (data not shown). We follow the same modelling protocol as for the case of CoHex^3+^ to derive effective potentials for this control model.

It may be noted that the D-D RDFs in the Mg^2+^ and CoHex^3+^ systems between the central beads of different DNA molecules (Figure 4A) appear very different. However, the final effective potentials for the D-D pair from both simulations (Figure 4D) have similar features. They are both slightly attractive from about 21 Å to the cut-off 25 Å. In the distance range below 21 Å, both effective potentials are repulsive. In the Mg^2+^ system, the RDF at distances below 25 Å has low amplitude since different DNA fragments repel each other due to electrostatic interactions. The presence of Mg^2+^ counterions cannot overcome this repulsion. In the more strongly coupled CoHex^3+^ system, the presence of the trivalent counterions creates an overall attraction between DNA segments that leads to DNA condensation. Therefore, it is expected that similar D-D potentials result in different D-D RDFs if different ions are present in the system.

Furthermore, the effective potentials for monovalent ion-ion interactions obtained under different conditions are virtually the same as illustrated in Figure 4F for the case of the Na-Cl pair. In Figure 4C the corresponding Na-Cl RDF shows, however, a small but noticeable difference between the aggregating (CoHex^3+^-system) and non-aggregating (Mg^2+^-system) simulations due to different average DNA-DNA distances. However, the final effective potentials from both systems are identical (Figure 4F). We attribute this to the fact that correlations between the different interaction terms are well captured in the present model. In fact, all ionic potentials for the same interaction types, but extracted from different underlying all-atom MD simulations for the present CG DNA model, are indistinguishable or very similar. As a further illustration, Supplementary Figure S5 compares ionic potentials extracted from three DNA all-atom MD simulations having different ionic and different DNA compositions. This behaviour implies good transferability of the derived potentials (see below for further discussion).

### DNA condensation in CG simulations

Having rigorously extracted effective potentials for the CG DNA model on the basis of the all-atom simulations, we next proceed to use them in large-scale simulations investigating DNA phase separation induced by the presence of explicit CoHex^3+^ ions. The applied CG approach treats long-range electrostatic interactions explicitly with the presence of all mobile ions in the system. The total solvent-mediated interaction potential between all charged sites in the system is a sum of a Coulombic potential scaled by the dielectric permittivity of water (ε=78), and a short-range non-bonded interaction (shown in the figures, e.g. Figure 4) within the cut-off distance, determined by the IMC procedure. This treatment of the long-range electrostatic interactions was validated in our previous work (54). Not only does this lead to a rigorous description of the important electrostatic interactions, but also enables the CG model to be used under varying ionic conditions. It is therefore possible to run CG simulations for other ion compositions and concentrations without having to rerun atomistic simulations. This means that in future studies, we can address salt dependent effects within the CG model. It may be noted that the SCG model (below) is not transferable as it does not include explicit ions. For other salt conditions, the effective potentials in the SCG model would need to be recalculated. This can be done within the CG model where we can easily change the ion composition since all electrostatics is explicit, while DNA-DNA potentials can, with a good confidence, be considered as independent of the salt content. So this two-step procedure is also practical if one wishes to generate SCG models for different ion conditions.

After the significant reduction in the number of degrees of freedom by the coarse-graining, we can easily simulate DNA condensation in a box of size of 150×150×150 nm^3^ for extended time. This is 1000 times larger volume compared to what is affordable for the all-atom MD simulations, which have used a box of 15×15×15 nm^3^. Here, 200 pieces of 100-bp CG DNA double helices are randomly placed in the box together with CoHex^3+^, potassium and sodium ions as well as the appropriate amount of chloride ions.

Figure 5A displays the short-range energy of the CG model as a function of time with representative snapshots and illustrates that DNA aggregation and phase separation occurs gradually during the simulation. The DNA condensate particle gets larger, forming a single fibre-like particle at the end of the simulation. Interestingly, the DNA molecules demonstrate short-ranged local order of hexagonal arrangement in the fibre bundle. This illustrates that the almost universal hexagonal packing of condensed DNA is intrinsically inherent in the chemical and physical properties of the DNA molecule as represented by the underlying all-atom force field and preserved in the effective potentials.

**Figure 5.**
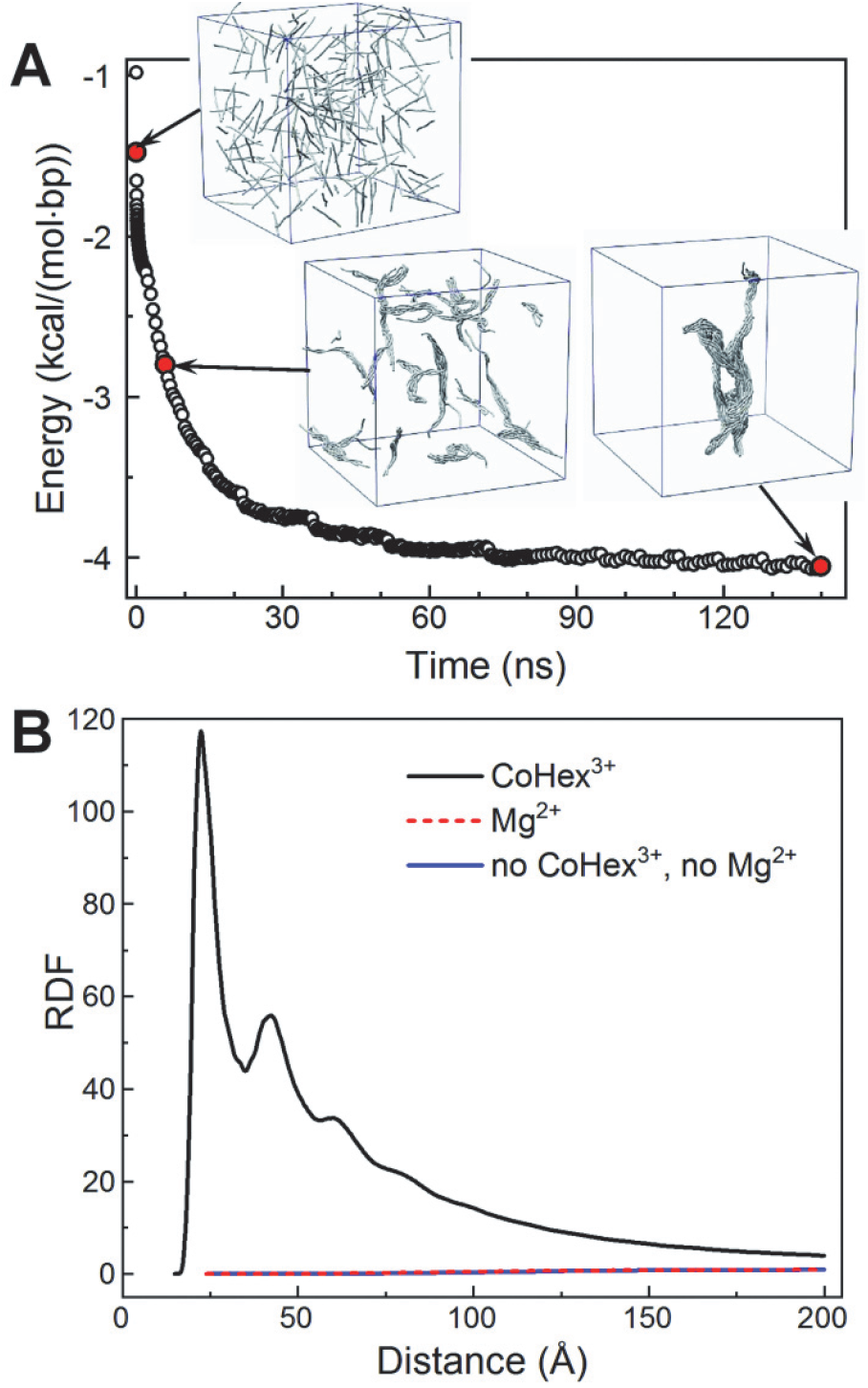
Illustration of DNA phase separation in CG simulations. (**A**) Snapshots and short-range interaction energy profile from the CG DNA simulation. Time points corresponding to each snapshot are indicated with red circles on the energy curve. DNA molecules are drawn as gray tubes following the path of the double helix; mobile ions are not shown. (**B**) Comparison of the D-D RDF determined in the CG simulations with the CoHex^3+^ (black), Mg^2+^ (red dashed) ions and with monovalent salt (blue).

The value of the shortest DNA-DNA distance is *r* = 22.5 Å, as it is exhibited by the first peak of the RDF in Figure 5B. This is in reasonable agreement with experimental data. Our own observations (unpublished data) from X-ray diffraction measurements of precipitated 177 bp DNA molecules, display a single broad Bragg peak at *q* = 0.26 Å^−1^. This corresponds to a lateral DNA-DNA separation in the range 24 Å – 27 Å (assuming hexagonal (*r* = *4π*/*q*√*3*) or lamellar (*r* = *2π*/*q*) packing). A shorter separation observed for bundled DNA in simulations compared to experiment is likely due to the CHARMM force field presently used in the underlying all-atom MD simulations (66).

To demonstrate the robustness of the CG DNA model, we conduct another simulation where the CoHex^3+^ ions are replaced by K^+^ and Na^+^ ions with equivalent amount of charge, using potentials obtained from the IMC procedure described above. The same CG box size and simulation protocol are adopted. The system exhibits no DNA condensation and the DNA-DNA interaction is repulsive as can be seen from the D-D RDF plotted in Figure 5B (blue line). In contrast to the system with CoHex^3+^, the amplitude of the D-D RDF is low in value, suggesting a large distance between DNA molecules. Additionally, Figure 5B displays the D-D RDF (dashed red line) obtained from a CG simulation where the CoHex^3+^ ions are replaced by Mg^2+^ ions, with all potentials obtained from the all-atom system comprising four DNA molecules and Mg^2+^ ions, as mentioned above in relation to Figure 4D-F. In agreement with the experimental data (1), this system is repulsive and phase separation does not occur. Hence, we can conclude that our CG DNA model is robust and produces realistic DNA aggregation behaviour in large-scale simulations in agreement with experiment.

The result illustrated in Figure 5B also lends support for the transferability of the effective ionic potentials obtained with the present approach. The CG simulation resulting in the repulsive D-D RDF (red line in Figure 5B) is performed for a system with 200 DNA molecules of 100 bp length containing only monovalent ions. In such a system the expected macroscopic behaviour is that DNA condensation should not occur. However, the effective ionic potentials used in the CG simulation resulting in this non-aggregating system are obtained from an all-atom MD simulation containing CoHex^3+^ ions that displays aggregation and bundling at all-atom level (Figure 2). As a further test of the transferability of the ionic potentials, we compare effective potentials for all monovalent ion interactions obtained by the IMC procedure from two different underlying all-atom MD simulations, displayed and discussed in Supplementary Figure S5. An additional test of transferability is illustrated in Supplementary Figure S6.

### Mesoscopic-scale simulations of DNA hexagonal phase separation

To investigate DNA phase separation at the mesoscopic level we performed one more step of coarse-graining, constructing the super coarse-grained (SCG) model. The excellent behaviour of the CG DNA model allowed us to confidently build a DNA model with even lower resolution and better performance in terms of computational time. The ions, as well as all electrostatic interactions are treated implicitly by the effective potentials. RDFs and effective potentials comprising the SCG DNA model are illustrated in Figure 4G-I. Due to the simplicity of the model there is no significant correlation among the three interaction terms; the bond and angle potential minima are at the same position as the maxima in the corresponding respective distributions. These two terms may in principle be modelled by simple harmonic functions. On the other hand, the non-bonded interaction term cannot be directly fitted by conventional functions, such as a Lennard-Jones or Debye-Hückel potentials. Specifically, although there is a dominant minimum at 23 Å in the non-bonded effective potential, which might be mimicked by a Lennard-Jones potential, the long-range behaviour of the IMC-computed potential is different, with a positive maximum at 35 Å followed by two relatively small minima at about 44 Å and 65 Å. Hence, the final effective potential contains interaction features that preserve the characteristics of the underlying fine-grained CG simulation as well as the all-atom MD simulation. The present systematic hierarchical multiscale modelling approach can thus preserve more detailed information even with a DNA representation as simple as beads-on-a-string.

The resulting SCG DNA model is used in mesoscale simulations of DNA phase separation for two types of systems. First, 400 relatively short DNA molecules, each one equivalent to 96 bp, are randomly placed in a 150×150×150 nm^3^ box. At the end of this simulation DNA condense into large particles consisting of more than 100 DNA molecules as illustrated in Figure 6A. Noticeably, these particles exhibit a hexagonally ordered structure resembling a liquid crystal, illustrated in the cross-section view in Figure 6B. Inside these aggregates, DNA molecules are arranged in such a way that the hexagonal structure can be seen from the cross-section of the particle, in agreement with experimental data obtained for short DNA molecules in the presence of the trivalent cations, CoHex^3+^ and spermidine^3+^ (18,19).

**Figure 6.**
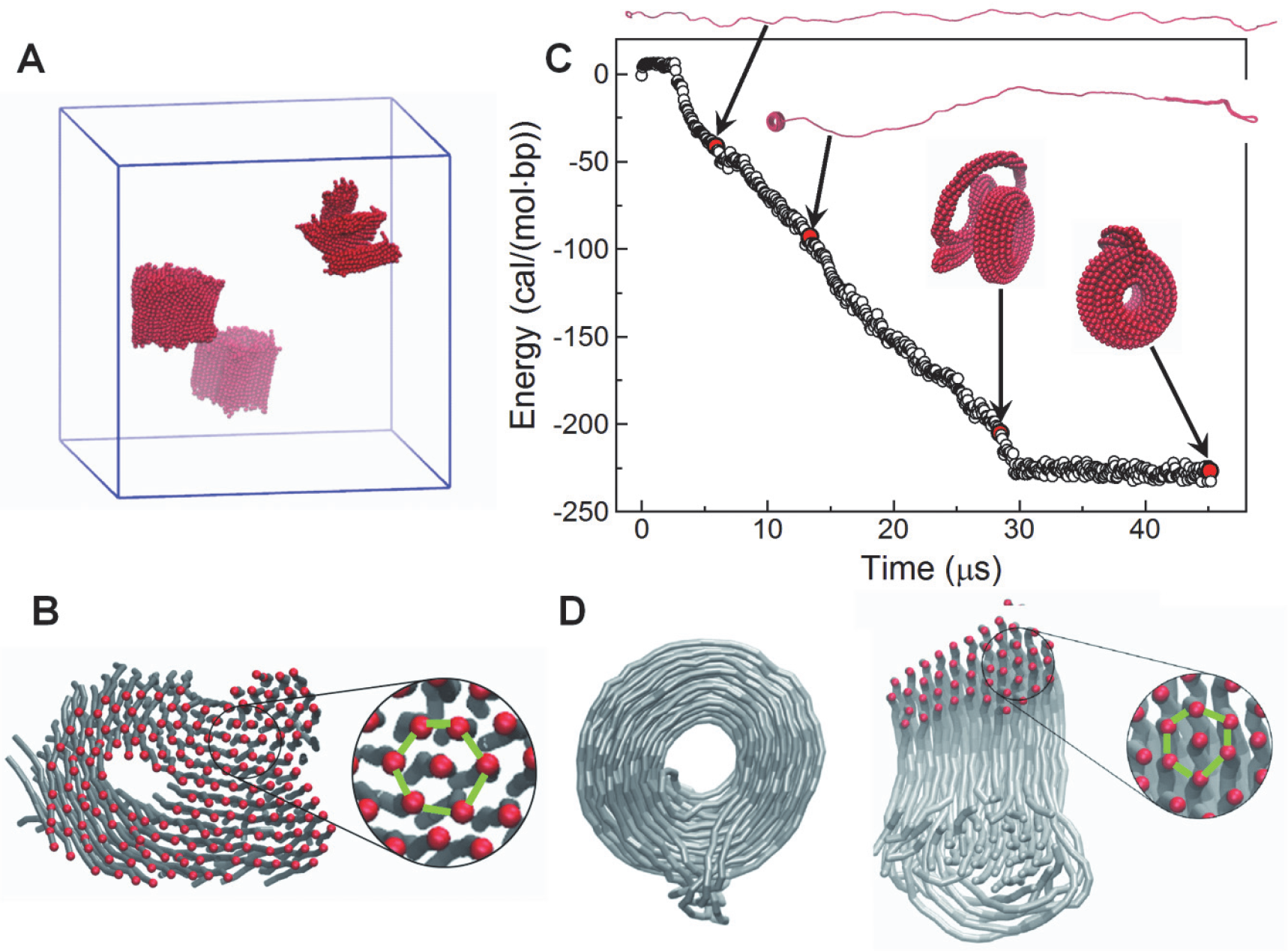
DNA aggregation and toroid formation simulated by the SCG DNA model. (**A**) Final configuration of DNA aggregation in a simulation with 400 DNA molecules. (**B**) Cross-section of one of the DNA condensed particle shown in (**A**). (**C-D**) Formation of toroidal structure in the SCG simulations of a 10.2 kbp DNA. (**C**) Energy profile and snapshots (normalized per DNA base pair) from one of the simulations. (**D**) Structure of the DNA toroid. Cartoon on the right-hand side shows cross-section through the toroid where the red dots illustrate DNA double helices near the cutting plane. The zoom-in illustrated in (**B**) and (**D**) show the DNA packing within the aggregates with green lines highlighting hexagonal arrangement of the DNA molecules.

Secondly, we perform simulations for a single 10.2 kb long DNA molecule with the SCG DNA model. The simulations start from a fully extended DNA conformation. After a short relaxation at the beginning, a loop or a bundle can form at either end of the DNA (see Supplementary Movie S1). The top snapshot in Figure 6C shows the DNA conformation after such a loop formation. Subsequently, the loop and bundle play the role of nucleation sites, attracting more DNA beads to form a toroid that grows in size (see Supplementary Movie S2). Towards the end of the simulation, the whole DNA molecule is condensed into one toroidal particle (see Supplementary Movie S3). The non-bonded interaction energy decreases as the toroid grows in size, illustrated by the energy profile in Figure 6C. Remarkably, as can be seen in Figure 6D, the cross section of the toroid shows that DNA is organized in well-defined hexagonal arrangement. These structural features are consistent with the reported electron microscopy studies (6).

Analysis of multiple simulations of single 10.2 kb long DNA molecules reveals that toroids are mainly formed in two ways. The first scenario suggests that a single loop at one end initiates the toroid formation, while the other end may eventually form a fibre that subsequently joins the toroid at the end of the simulation (see Supplementary movies S1, S2 and S3). Secondly, a loop may get formed at each end in the beginning of the simulation with the simultaneous growth of two toroids that eventually join together (exemplified in Supplementary movie S4). An interesting feature of DNA toroid formation and size increase, is the sliding motion between contacting DNA segments. DNA toroids can adjust their conformation through this motion simultaneously with the growth due to rolling and attracting more DNA fragments. It should be noted that not only toroids are formed, but the final condensed structure can be fibre-like, which has also been observed in EM experiments (22). In the simulations, the toroid shape is, however, somewhat more frequently occurring. We also simulate a lambda-DNA size single DNA molecules (48 kbp) that also form toroids by the mechanism with nucleation loops at both ends (data not shown).

Next we perform an analysis of the mechanism of DNA condensation that results in toroid or fibre bundle formation. We pay specific attention to the pathways and the initial events during DNA condensation that leads to formation of the major final states of toroid or fibre-like morphologies. To this end we perform 67 independent SCG simulations of the 10.2 kbp DNA (having same initial coordinates but with different starting velocities). Although the mechanism of DNA condensation to either toroid or fibre is complex and stochastic, it is possible to identify several intermediate states as well as transitions between them. The formation of either toroid or fibre can be divided into two stages. The first stage can be characterized as nucleation. The second stage includes the growth of the nucleation site that occurs by pulling in more and more DNA chains, and by the sliding mechanism mentioned above. Figure 7 summarizes the early event intermediate states and transitions between them. Supplementary Table S3 gives a description of those frequent initial transient structures. In the first few nanoseconds of each simulation, several transient states can be observed, such as state **b** and state **k** in the figure. These are usually simple structures with short lifetimes. As the simulation proceeds, more stable structures can be formed, for example, bundles consisting of three DNA chains (state **c** in the figure) and the double DNA chain loop (state **g**). Based on these more stable structures, the “growth” mechanism can be effective with the remaining part of the DNA chain being pulled in to form larger and more stable structures.

**Figure 7.**
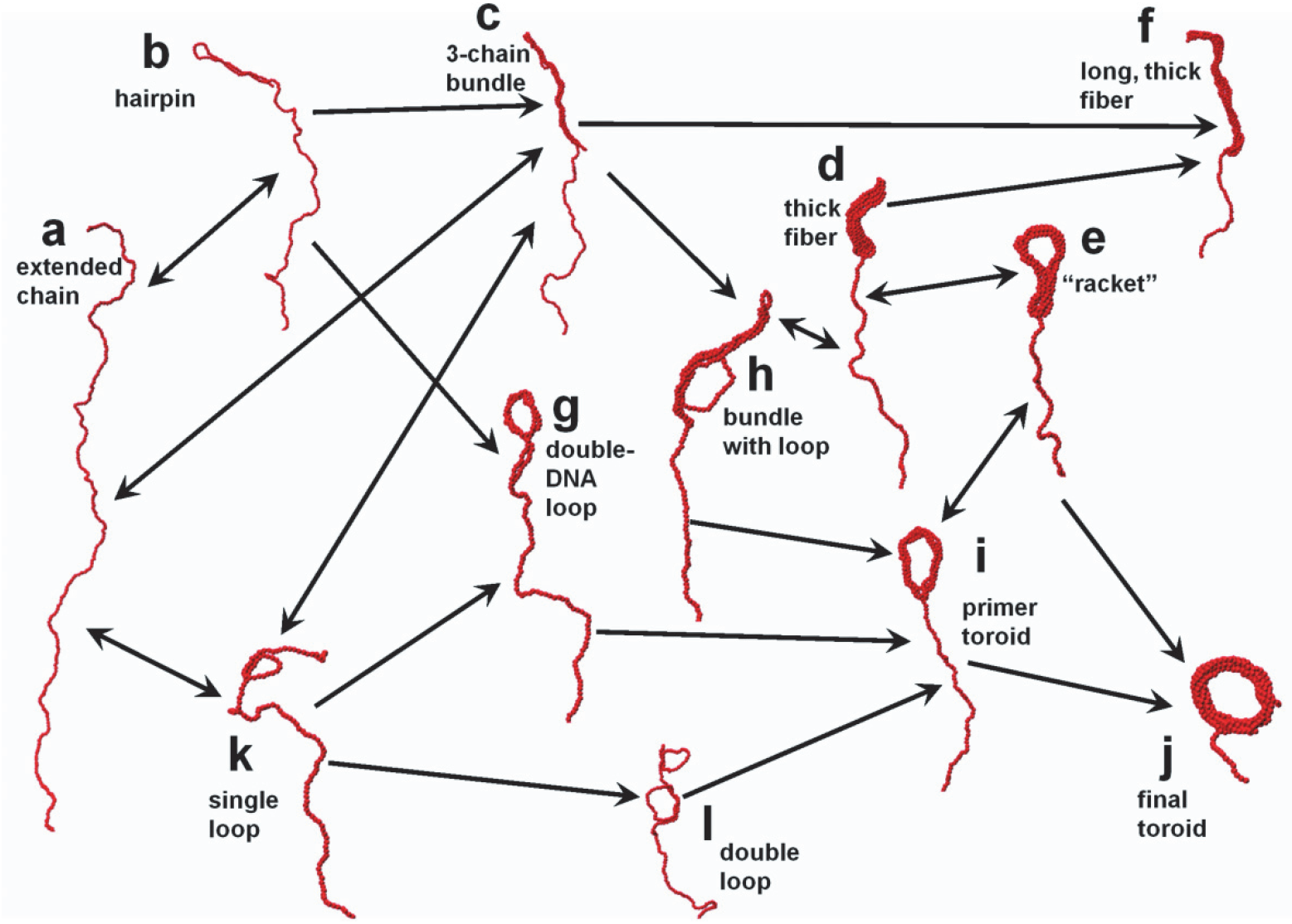
Pathways and intermediate states during formation of DNA toroid and fibre. The structural features of each state are described in more details in Supplementary Table S3. Arrows indicate the transition directions among the states, as observed in the SCG simulations.

In the early stage of the simulation, fibre-like structures and toroid-like structures can convert between each other. These structures are usually no more than three DNA chains thick. Hence, the energy barrier involved in these transitions should not be prohibitive. For example, a three DNA chain bundle (state **c**) can convert to a single DNA chain loop (state **h**) to form a toroid-like structure (state **i**). On the other hand, this toroid-like structure (state **i**) is soft enough, so that it can go to a bundle (state **d**) just by closing the hole in the middle. These transitions between fibre-like and toroid-like states rarely happen in the subsequent stages of the simulation when more DNA chains become condensed.

We measure the dimensions of toroids formed by the 10kb DNA in multiple simulations shown in Supplementary Figure S7. The details of the calculations are given in Supplementary Figure S8. In brief, the projected toroid density map was measured using two circles, one to fit the outer perimeter of the toroid and one to fit the hole. The toroid diameter is an arithmetic average of the diameters of the two circles. Toroid thickness is the difference of the radii of the two circles (see Figure S7). The average thickness of the toroid is about 12 nm, which is smaller than in experiments with 3 kbp DNA (~25 nm) (22). Similarly, the observed diameter (~22 nm) and hole diameters (~10 nm) are also smaller than the dimensions observed in the experiments (22). These values are reasonable given the differences between experiments and simulations. The reason for smaller diameter and thickness obtained in simulations comparing to the experiments, might be due to the fact that the simulations contain only a single DNA molecule. Experimentally the toroids formed from DNA of sizes below 50 kbp contain several DNA molecules that leads to larger and thicker toroids (22). On the other hand, the diameter of the hole is expected to depend on nucleation loop size mechanism of toroid formation (22) and on the DNA bending properties as well as on effective DNA-DNA attraction. The small hole diameter observed in the simulations is most certainly caused by the intrinsic properties of the underlying force field, which compared to real DNA may represent a mechanically more flexible DNA with stronger attraction leading to shorter DNA-DNA distances and tighter packing in the toroid. The fact that toroid dimensions are strongly affected by electrostatic interactions between DNA molecules was earlier shown by Hud and co-workers (21), who demonstrated a pronounced dependence of toroid diameter and thickness on ionic conditions.

## CONCLUSIONS

In conclusion, we have used a rigorous hierarchical multiscale simulation scheme, which enables simulation of DNA condensation at mesoscale levels. The phenomenon of DNA condensation induced by multivalent ions is clearly inherent in the chemical properties of the DNA molecule. Inspired by this fact, we reasoned that a chemically based starting point, using state-of-the-art molecular force fields for all-atom MD simulations, followed by systematic coarse-graining, and using the IMC approach to extract solvent mediated effective CG potentials, would preserve those features of DNA in the CG models. Indeed, DNA condensation induced by the three-valent cobalt(III)-hexammine ions was demonstrated in large-scale simulations of hundreds of DNA molecules, which exhibited correct experimental structural features. We used a hierarchical approach where the CG model was further coarse-grained to a “super CG” model. Simulations at mesoscale level (10.2 kbp DNA) demonstrated toroid formation into hexagonally packed DNA, with reasonable dimensions in qualitative agreement with experimental observations. These results were obtained without any other underlying assumptions except for the all-atom force field and the DNA topology model adopted in the CG simulations and used no adjustable parameters.

The present parameterisation of the CG DNA model is not sequence specific, although this can in principle be introduced at the two-bp step level in the mapping of the all atom simulations. This would however, considerably increase the dimensionality of the CG as all 10 possible unique two-bp steps would have to be parameterised individually and described by separate effective potentials in the IMC procedure. Here we consider a general DNA sequence with mixed AT/GC content close to 50% in the 4-DNA all-atom simulations and 42% GC in the single DNA all-atom simulations. We compare with generic experiments on salt dependence of DNA persistence length and DNA condensation where DNA sequence specificity has generally not been considered. Hence, this approximation is justified. But it should be mentioned that various DNA breathing and bubbling events (bp unpairing) can happen and are allowed in the all-atom simulations and will then be translated to the effective potentials, which in turn means such events are taken into account in the CG model in an average way and would affect the persistence length. Based on our analysis, the frequency of these are force field dependent but effects on the persistence length are minor (Minhas et al, to be published).

In the present work we used all-atom MD simulations based on the CHARMM27 force field. However, we recently demonstrated similar behaviour in all-atom simulations that showed DNA-DNA attraction and bundling using both CHARMM36 and AMBER parameters (51). It should furthermore be of interest to investigate how the mesoscale simulation results depend on the CG topology comparing different CG DNA models, which include DNA sequence specificity (37,47). It may be noted that the present CG DNA model is not sequence specific, but such an extension can be implemented in our model (53).

The present successful approach lends support for developing CG models for more complicated systems exhibiting DNA compaction at mesoscale level such as chromatin and individual chromosomes. Such models will help understanding the compaction behaviour of chromatin as a function of various variables known to regulate genome compaction such as histone tail modifications that change electrostatic interactions. Multiscale modelling of nucleosomes and chromatin fibres, following the present approach requires development of a corresponding CG model describing histones and their interactions. The model shall take into account important hydrophobic and hydrogen bonding effects, which can be difficult to describe within such CG models and that may require the combination with trained top-down approaches for parameterisation. Hence, this is still a major step to be accomplished but our present work along those lines shows that such an extension may be feasible (A. Mirzoev et al, unpublished). Finally, in order to rigorously evaluate the time dynamics in the mesoscale simulations, generalized Langevin dynamics with friction parameters extracted from underlying detailed simulations can be performed, which enables the study of time-dependent condensation behaviour (67,68).

## DATA AVAILABILITY

All data needed to evaluate the conclusions in the paper are present in the paper and/or the Supplementary Data. Additional data related to this paper may be requested from the authors.

## SUPPLEMENTARY DATA

Supplementary Data are available at NAR online.

## ACKNOWLEDGEMENT

We are indebted to Profs Aatto Laaksonen and Francesca Mocci for discussions and suggestions. We acknowledge the generous support of computer time allocation from the National Supercomputing Centre (NSCC) Singapore.

## FUNDING

This work was supported by the Singapore Ministry of Education Academic Research Fund (AcRF) Tier 2 (MOE2014-T2-1-123 (ARC51/14)) and Tier 3 (MOE2012-T3-1-001) grants (to L.N.) and by the Swedish Research Council (grant 2017-03950 to A.P.L.).

## CONFLICT OF INTEREST

The authors declare that they have no conflict of interest.

## Supplementary Data for the manuscript

### SUPPLEMENTARY METHODS

#### A. All-atom Molecular Dynamics simulations

In the present work, we use the atomistic CHARMM27 force field (1,2) as a high-resolution reference in the inverse Monte Carlo (IMC) procedure to extract effective potentials for the coarse-grained (CG) model. In previous work (3) we have also tested the CHARMM36 and AMBER bsc0 force fields, which showed similar results concerning DNA aggregation in the presence of CoHex^3+^. Of the force fields tested, CHARMM27 provides better preservation of DNA B-form internal structure. Moreover, it has produced an excellent agreement with experimental DNA persistence length data. (see main text and (4)). The all-atom MD simulation is set up with four 36 bp-long double helix DNA molecules placed in a cubic simulation cell with periodic boundaries. The DNA sequences are the same as in our previous work (3) representing a 50-50% mixture of AT and CG pairs as shown below.

5’-ATTAATGGAACGTAGCATATTCTTCAAGTTGTCACG-3’

5’-CAAAACCTGATGCACACTGTAACATGAGATCCCGCG-3’

5’-TCGGCTTATAGAGGGCCAGCTCGTATCGACGGACCG-3’

5’-GCTAGTACCCCACCAATTTAGGCGAAAGGAGTCTGC-3’

Improved force field parameters for the CoHex^3+^ ions were derived in our recent study (3). The number of CoHex^3+^ ions is determined in such way that the charge carried by CoHex^3+^ is 1.5 times the charge of DNA, which should ensure attraction between DNA molecules. Additional salt is added to reach a salt concentration of 50 mM K^+^ and 35 mM Na^+^, with neutralizing amount of Cl^−^ ions. The improved ion parameters by Yoo and Aksimentiev (5) are used throughout all simulations for all ions except CoHex^3+^ and the TIP3P water model is utilized. The system is illustrated in Figure 2 of the main text.

In total, three trajectories of 1 μs each are generated from the same starting configuration with DNA oligomers placed randomly in the simulation cell and different starting velocities. The system was equilibrated in three stages. First, DNA and CoHex^3+^ ions are restrained while the system reaches a target temperature of 298 K. Then pressure coupling is turned on to maintain 1.013 bar pressure with only DNA molecules restrained, after which the unrestrained equilibration is conducted under constant temperature and constant pressure for 500 ns.

The production phase is conducted for at least 500 ns. The velocity rescale thermostat (6) and Parrinello-Rahman barostat (7,8) regulate respectively temperature and pressure. Bonds are constrained with the LINCS algorithm (9) implemented in GROMACS 5.1 (10), which enables a 2 fs time step. Electrostatic interaction is treated with particle mesh Ewald method (11) with 10 Å real space cut-off. The van der Waals interaction is treated with a cut-off scheme with the potential shifted to zero at cut-off distance of 10 Å.

Also, a control experiment is set up with the same all-atom simulation box substituting all CoHex^3+^ ions for the same number of Mg^2+^ ions. Additional Na^+^ and K^+^ ions are added to keep charge neutrality. For validation of the CG DNA model by persistence length calculation, a single 40 base-pair DNA with sequence 5’-GGATTAATGGAACGTAGCATATTCTTCAAGTTGTCACGCC-3’ (42.5% GC content) was used. To mimic typical experimental conditions the NaCl concentration is set to 130 mM. All other conditions are similar to the aforementioned simulations with CoHex^3+^.

#### B. Coarse-grained DNA models

In order to reach the mesoscale level of DNA modelling, we have performed coarse-graining at two spatial scales resulting in two coarse-grained DNA models with different resolutions. The model at the first level is mapped from all-atom DNA (Figure 1A), and is called the CG DNA model (Figure 1B). The second model with lower resolution is called the super CG DNA (SCG) model (Figure 1C). The CG DNA model is designed according to the same concepts as our previous model, which was shown to well reproduce DNA persistence length dependence on salt concentration over a wide concentration range (4). It is simple enough in its design and yet captures the structural form and properties of double helical B-form DNA. The DNA is modelled with consecutive units of five beads, representing two base pairs of DNA (Figure 1D). Four beads represent the phosphate groups (denoted “P”), while the central one (denoted “D”) is assigned for the four nucleosides. There are totally four types of bonds and three types of angles in the bonded interaction terms (Figure 1D), which maintain a helical B-DNA structure where two strands of the phosphate groups form the major and minor grooves (Figure 1B). The pairwise bonds are D-D, D-P, P-P along the strand and P-P cross-strand defined as a shortest P-P bond over the minor groove. The angle bonds are D-D-D, P-D-P, and P-P-P. The units at each end of the DNA double helices are treated specially. Since these units have the opportunity of closely contacting solvent and ions, we used dedicated bead types (denoted “DT” and “PT”) to allow different non-bonded interactions. However, the bonded interaction terms remain the same as the non-terminal beads.

The model, which includes explicit mobile ions and explicit charges of the DNA phosphate groups, pays particular emphasis to electrostatic interactions, where every P-bead has charge −1e, and D-beads are kept neutral. The solvent is considered implicitly, via use of the effective short-range potentials (computed by the IMC method) and screening the electrostatic interactions according to the solvent dielectric permittivity ε =78.

Despite looking similar to its ancestor (4), the present model has several very important and considerably different features. First, the underlying all-atom MD simulations are significantly longer (an order of magnitude compared to the previous model). Second, all the effective potentials are derived from the underlying all-atom MD simulation using the Inverse Monte Carlo method. Particularly, all non-bonded interactions are not simply a combination of electrostatic and repulsive truncated Lennard-Jones (as in the case of ref (4)), but are rather complex functions which are capable of describing distribution of ions around DNA in solution according to the underlying atomistic simulations. In the present model, both bonded and non-bonded potentials (shown in Figure S2) are tabulated and are not fitted to any functional form and contain no adjustable parameters.

In the CG model, bonded interactions are capable of maintaining the structure as well as the mechanical property of DNA. But to reduce the correlation between bonded and non-bonded interaction terms, we excluded short-range effective interactions between non-bonded pairs within the same duplex. Conventional treatment of excluding electrostatic interactions between sites with explicit bond or angle is applied in all cases.

The next level, the beads-on-string SCG model of the DNA prioritizes computation performance over complexity (Figure 1C). In the SCG model, single type of uncharged beads (called ”S”) represents three units of the CG DNA model that corresponds to six DNA base pairs. Bonded interaction terms include one bond and one angle potentials. The non-bonded interactions are treated in conventional manner and include the intermolecular interaction between the S beads as well as the intramolecular terms excluding “1-2” and “1-3” pairs. Compared to the CG DNA model, in the SCG model not only the solvent is considered implicitly, but electrostatic interactions are implicitly included into the effective potentials. The effective potentials are derived by the IMC procedure after mapping the CG-model trajectory to the SCG scheme.

#### C. Inverse Monte Carlo procedure for extraction of effective CG potentials

In the CG DNA model, all effective interaction potentials are determined solely by IMC. The only input information is the structural properties extracted from all-atom MD simulations in terms of the relevant RDFs between the sites of the CG model. These are obtained by mapping the all-atom MD trajectory from the four-DNA system to a corresponding CG site trajectory of the MD simulation. Thus, no empirical parameters are used in the CG DNA model and it rests solely on the parameters from the all-atom CHARMM27 DNA as well on the CPMD optimized CoHex^3+^ (3).

To derive an effective potential function, the IMC procedure uses a radial distribution function (RDF) corresponding to each interaction in the CG system. During mapping of the atomistic trajectory to a CG trajectory, the “P” sites of the CG DNA model are defined as centre-of-masses of the corresponding phosphate groups (−PO_4_^−^)- of the DNA backbone. The “D” sites are determined as centre of masses of the four nucleosides constituting the corresponding two base pair fragment. To avoid end-effect ”contamination” of the distribution statistics, the ends of the DNA double helices are treated as separate types (named DT and PT). Thus the total number of bead types in the system is eight: comprising four DNA beads (D, P, as well as terminal DT, PT), one of CoHex^3+^, one of K^+^, one Na^+^ and one Cl^−^; which gives a total number 36 non-bonded interaction terms. All-atom MD trajectories from the three simulations were converted into CG trajectories by the mapping procedure described above. Convergence of in DNA aggregation is confirmed by comparing DNA-DNA RDFs from trajectory segments at different simulation times. Each final set of RDF is calculated as an average over the equilibrated sections of all three independent trajectories with a total length of 1.5 μs.

The IMC calculation is carried out with the MagiC software (12,13) (which is also used for bead-mapping, RDF calculation, analysis and export of the resulting potentials). A zero potential is used as the first trial potential for non-bonded interactions, while a potential of mean force (defined as *U_PMF_* = –*k_B_T ln*(*g(r)*) is applied as a trial for bond and angle interactions. The effective potentials are refined in about 20 IMC iterations, with 100 parallel Monte Carlo sampling simulations in each iteration. In each sampling thread, 300 million Monte Carlo steps are performed with acceptance ratio maintained at about 50%. The first half of each thread is considered as equilibration. Potentials for the CG DNA model in presence of CoHex^3+^ can be found in Supplementary Figure S3. All charge sites interact by a Coulombic potential (scaled by the dielectric constant) and treated by the Ewald summation. This means that DNA interacts by this potential beyond 25 Å, which is the cut-off used only for the short-range part of the total potential. Long-range electrostatic interactions are treated using Ewald summation with real space cut-off being 40 Å. The dielectric constant is set to 78.0. Monte Carlo sampling is performed within a constant volume and constant temperature ensemble for the same number of DNA (of the same length) and ions, and in the box of the same size as in the underlying atomistic simulations. The procedure described above is followed in a similar way for deriving the effective CG potentials from the other all-atom systems, namely the one with four DNA and Mg^2+^ ions and the one comprising a single DNA and monovalent salt.

A table summarising the different system setups and CPU performance is provided below (Table S1) and a list of all effective potentials and underlining all-atom MD simulations used to derive them is given in Table S2 located at the end of this section.

The effective potentials of the SCG model are derived in the same way as for the CG model applying the IMC method. Specifically, the trajectories from the MD simulation of the CG DNA systems are mapped to a SCG trajectory and from these data a bond distribution (S-S) and an angle distribution (S-S-S) as well as one non-bonded DNA-DNA RDF (S-S) were calculated with 200 Å cut off. The IMC calculations are started from zero potential for non-bonded interaction, and with potential of mean force for bond and angle interactions. Temperature and volume are kept constant during sampling with 100 parallel Monte Carlo simulations. Typically, the effective potentials converge in about 25 iterations.

#### D. Coarse-grained simulations in the CG model

The tabulated interaction potentials for the CG DNA model, obtained as described above are used in the CG MD simulations with a significantly bigger simulation box compared to the all-atom simulations. The simulations with the CG and SCG DNA models are conducted using the LAMMPS package (14) with the NVT ensemble. For the CG simulations, 200 pieces of 100-bp DNA (50 CG-DNA units) are randomly placed in a 150×150×150 nm^3^ simulation box. The box also contains CoHex^3+^ ions (carrying charge twice exceeding the one on the DNA) with 10 mM K^+^ and 10 mM Na^+^ ions as well as the neutralizing number of Cl^−^ ions. The CG simulation is started with a 1 fs time step to reach a stable temperature before switching to a 2 fs time step for next 1 million steps of equilibration. Langevin dynamics with damping parameter being 10 ps, is used to initiate the simulation. Finally, the production simulation is performed with 5 fs time step with a velocity rescale algorithm regulating system temperature. The particle-particle particle-mesh (PPPM) method is used to calculate electrostatic energies with a 40 Å real space cut-off. The same cut-off is applied for the short-range interactions. This procedure for CG simulations is conducted analogously for the systems with Mg^2+^ ions and with a single DNA at varying monovalent salt concentrations (persistence length validation). As noted above the short-range effective interactions within the same duplex are excluded. However, it is possible that DNA chain can be bent and have head-tail contact in simulations with long DNA. In this case, the short-range contacts should be treated in the same way as inter-DNA interaction. In the current CGMD simulations, this is implemented by introducing virtual bonds along the DNA chain. Specifically, on the central D-D chain, 1-3, 1-4, 1-5 and 1-6 D-D beads are connected by bonds with zero interaction potential. By excluding all 1-4 interaction pairs, all bead pairs within 6 units (12 bp) are excluded from the short-range interaction. Note that this exclusion rule together with the short-range cut-off effectively eliminates all pairs in short segments along the DNA duplex while keeping those pairs far away along DNA but spatially close.

#### E. Coarse-grained simulations using the SCG DNA model

Within the SCG model, the electrostatic interactions are treated implicitly since they are effectively included into the SCG effective potentials. The simulation is started with a randomly generated velocity at 298 K. To stabilize the temperature at 298 K, velocity rescaling is used during the 10^5^ equilibration 5 fs-long steps. For the production stages, the time step is set to 200 fs with damping parameter of the Langevin thermostat at 298 K is set to 100 ps to facilitate fast sampling. Two sets of the SCG simulations are carried out: with multiply short DNA (400 pieces of 96 bp DNA, represented by a chain of 16 S-beads); and for single 10.2 kbp-long DNA (1700 S-beads). For multiply short DNAs simulation box was set to 150×150×150 nm^3^ and production run consists of 4×10^7^ steps; for single long DNA the box is 3450×3450×3450 nm^3^ and data collected for 2×10^8^ steps.

#### F. Persistence length calculation

We perform all-atom MD simulations of a single 40 bp DNA in the presence of physiological salt (130 mM NaCl). We perform the mapping of this trajectory to the CG model and extract effective IMC potentials (Figure S1 shows all the effective potentials and they are listed in Table S1). There is no DNA-DNA short-range effective pair interaction applied in the simulations. However, do note that this is consistent with our CG model with the CoHex^3+^ case even with DNA as long as 500 bp. With single DNA and monovalent salt, the electrostatic interaction results in no close contact (shorter than the short-range cut-off) between DNA beads.

A 500 bp long CG DNA is then simulated in a range of NaCl salt concentrations from 0.1 mM to 100 mM. All the simulations are run for at least 3 μs and employ a protocol described above. The values of persistence length, *L_p_*, are calculated by equation: *L_p_* = −*L_c_*/(ln<*cosα*>), where *L_c_* is the contour length of the DNA and <cos*α*> is the average of the cosine between two adjacent segments over the length of the simulation for the corresponding contour length. A 500 bp DNA is divided into 5 equal length segments (100 bp, *L_c_* ≈ 330 Å). That results in 5 data points for each salt concentration. The values of *L_p_* are determined from the slope of the plot *L_c_* versus ln<cos*α*> (4). The standard deviation of *L_p_* is evaluated with these 5 data points. The *L_p_*. values calculated from different parts of the trajectories show similar values, which confirms convergence of the simulations. Our simulation results are compared with experimental data and with the data of other simulation studies.

---

**Table S1.** Interaction potentials for the coarse-grained DNA model derived in this work. “✓” indicates the presence of the interaction term in the model represented by the column. The DNA-DNA interaction terms are defined as inter-DNA interactions. The short-range intramolecular DNA bead-bead interactions within the same duplex are excluded while distant interaction pairs along the DNA helix, but spatially close in space, are included. This is implemented by excluding pairs within 6 units of DNA using virtual bonds. In the case of a single DNA in monovalent ion solution, all intra-DNA pairs are excluded by setting all DNA-DNA pair potentials to zero since in 1:1 salt electrostatic repulsion keeps DNA segments beyond the cut-off of short-range effective interactions.

| Type of potential | Interaction term | All atom simulation used to parameterise a given CG model |  |  |
| --- | --- | --- | --- | --- |
|  |  | 4DNA+CoHex<sup>3+</sup> | 4DNA + Mg<sup>2+</sup> | 1 DNA |
| Bonds | D-D | ✓ | ✓ | ✓ |
|  | D-P | ✓ | ✓ | ✓ |
|  | P-P (backbone) | ✓ | ✓ | ✓ |
|  | P-P (cross minor groove) | ✓ | ✓ | ✓ |
| Angles | P-D-P | ✓ | ✓ | ✓ |
|  | P-P-P | ✓ | ✓ | ✓ |
|  | D-D-D | ✓ | ✓ | ✓ |
| Non-bonded potentials | DT-DT* | ✓ | ✓ |  |
|  | DT-PT* | ✓ | ✓ |  |
|  | DT-D* | ✓ | ✓ |  |
|  | DT-P* | ✓ | ✓ |  |
|  | DT-Co | ✓ |  |  |
|  | DT-K | ✓ | ✓ |  |
|  | DT-Na | ✓ | ✓ |  |
|  | DT-Cl | ✓ | ✓ |  |
|  | PT-PT* | ✓ | ✓ |  |
|  | PT-D* | ✓ | ✓ |  |
|  | PT-P* | ✓ | ✓ |  |
|  | PT-Co | ✓ |  |  |
|  | PT-K | ✓ | ✓ |  |
|  | PT-Na | ✓ | ✓ |  |
|  | PT-Cl | ✓ | ✓ |  |
|  | D-D* | ✓ | ✓ |  |
|  | D-P* | ✓ | ✓ |  |
|  | D-Co | ✓ |  |  |
|  | D-K | ✓ | ✓ |  |
|  | D-Na | ✓ | ✓ | ✓ |
|  | D-Cl | ✓ | ✓ | ✓ |
|  | P-P* | ✓ | ✓ |  |
|  | P-Co | ✓ |  |  |
|  | P-K | ✓ | ✓ |  |
|  | P-Na | ✓ | ✓ | ✓ |
|  | P-Cl | ✓ | ✓ | ✓ |
|  | Co-Co | ✓ |  |  |
|  | Co-K | ✓ |  |  |
|  | Co-Na | ✓ |  |  |
|  | Co-Cl | ✓ |  |  |
|  | K-K | ✓ | ✓ |  |
|  | K-Na | ✓ | ✓ |  |
|  | K-Cl | ✓ | ✓ |  |
|  | Na-Na | ✓ | ✓ | ✓ |
|  | Na-Cl | ✓ | ✓ | ✓ |
|  | Cl-Cl | ✓ | ✓ | ✓ |
|  | DT-Mg |  | ✓ |  |
|  | PT-Mg |  | ✓ |  |
|  | D-Mg |  | ✓ |  |
|  | P-MG |  | ✓ |  |
|  | Mg-Mg |  | ✓ |  |
|  | Mg-K |  | ✓ |  |
|  | Mg-Na |  | ✓ |  |
|  | Mg-Cl |  | ✓ |  |
| <sup>1</sup>Electrostatic Interaction | all non-bonded pairs | ✓ | ✓ | ✓ |

<sup>1</sup> It should be noted that we do not directly parameterise electrostatic interactions, as they are defined by bead-charges and the solvent dielectric permittivity. Since the charges and the solvent dielectric permittivity are set in the same way for the aforementioned models, the electrostatic interactions are shared by these CG models.

**Table S2.** Comparison of system size and CPU usage in simulations with all-atom, CG and SCG model. Data for the CG model is estimated after DNA aggregation, which is slower than a homogeneous (dispersed) system, due to CPU load imbalance. Performance can be improved by a better load distribution algorithm. Data for the SCG model is taken from the single DNA simulation.

| Model | Box size (nm) | Number of atom/beads | DNA length (bp) | Time step (fs) | CPU time per 100 ns (hr) | Software package |
| --- | --- | --- | --- | --- | --- | --- |
| All-atom | 16.6 | 456,097 | 36 | 2 | 17831.54 | GROMACS |
| CG | 150.0 | 142,632 | 96 | 5 | 27783.32 | LAMMPS |
| SCG | 3450.0 | 1700 | 10,200 | 200 | 1.68 | LAMMPS |

## SUPPLEMENTARY DATA

**Figure S1. RDFs and potentials used for the calculation of persistence length**

Interaction potentials calculated by the IMC procedure using RDFs obtained in an all-atom MD simulation of one single 40 bp DNA in the presence of 130 mM NaCl were used for validation of the CG DNA model. These potentials were used to determine salt dependence of the DNA persistence length (see Figure 3 of the main text).

**Figure S1.**
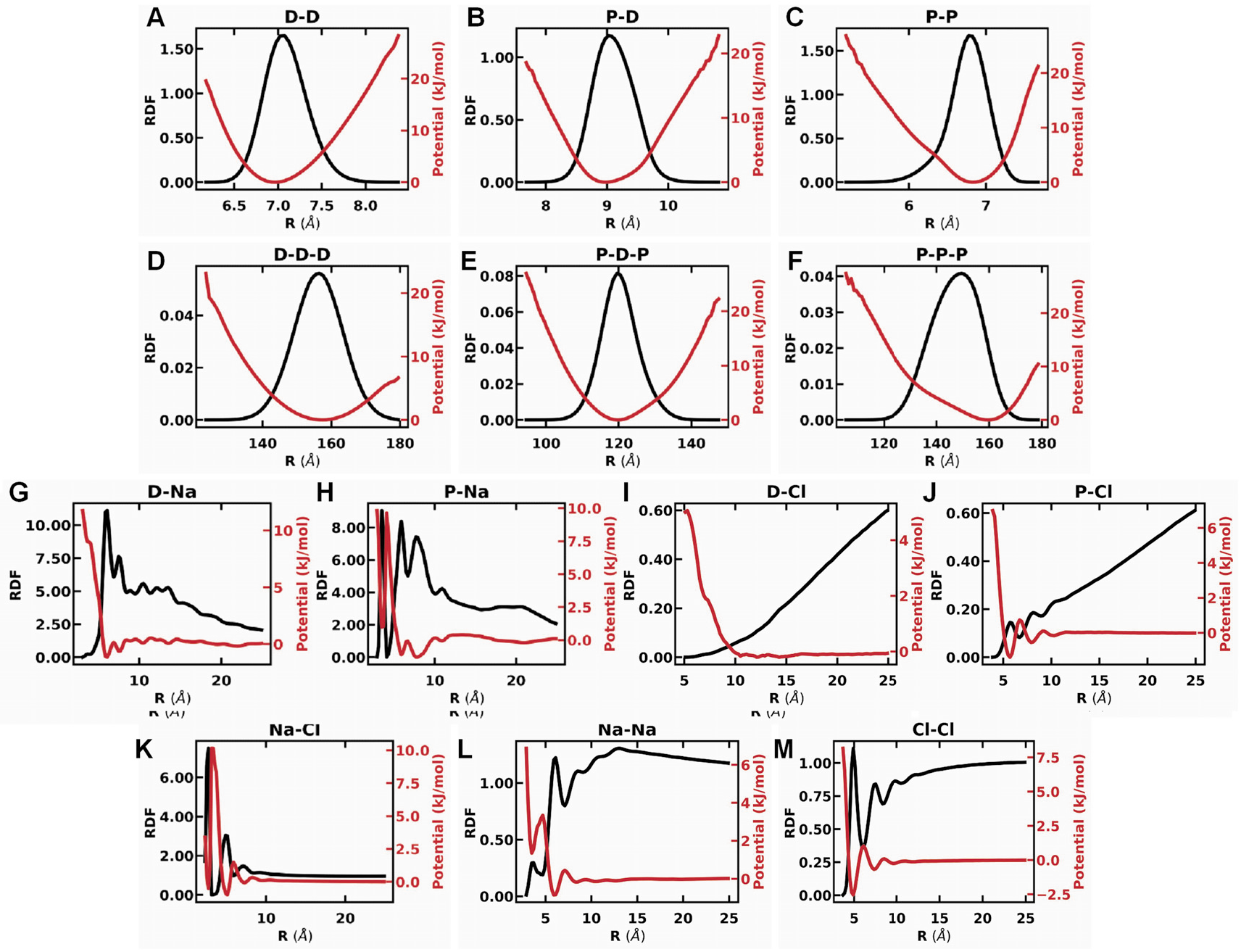
RDFs (black) and effective potentials (red) for interaction terms in the CG DNA model calculated from all-atom MD simulation of DNA in monovalent salt. Panels **A-C** and **D-F** show data for respectively CG DNA bonds and bonded angles; panels G-J are for the DNA – ion interactions; panels K-M are for the ion – ion RDFs and potentials. Identity of the interaction terms is shown at the top of the graphs.

**Figure S2. Snapshots from different all-atom and CG MD simulations modelling DNA-DNA interaction in systems with CoHex^3+^ cations.**

Figure S2 (A-C) illustrates variability of DNA bundling from snapshots in the three all-atom MD simulations. The simpler energy profile and significantly longer sampling in the CG MD simulation results in minor variability between different simulations (Figure S2D). Although all three simulations are technically converged (by standard criteria such as constant total energy and their components), when aggregation occurs the DNA bundles may be trapped in metastable states, which is unavoidable in any all-atom MD simulation of a complex system. This is why we perform three independent simulations and used the average RDFs.

**Figure S2.**
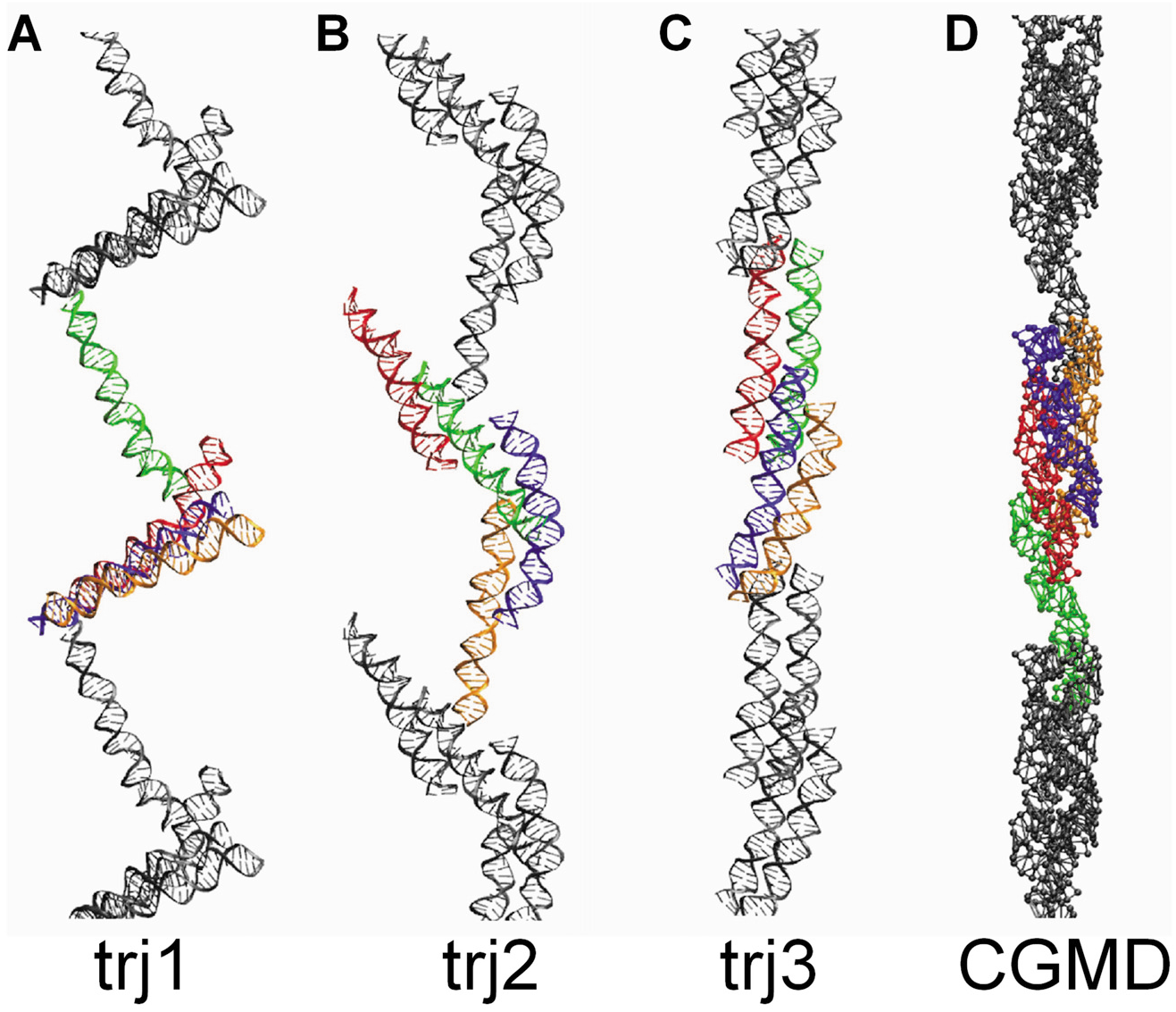
(**A-C**) Final snapshot conformations obtained for all-atom trajectories in the MD simulations with CoHex^3+^. DNA double helices are coloured differently. Periodic images of the DNAs are coloured grey. (**D**) A snapshot of the final conformation in a coarse-grained MD simulation using conditions similar to the all-atom systems and interaction potentials determined from the simulations shown in (**A-C**).

**Figure S3. The complete set of RDF and converged effective potentials of the CG model.**

Below, all the RDFs and effective potentials corresponding to all interaction terms in the CG-model simulations with CoHex^3+^ are displayed. DNA is modelled by four bead types (DT, PT, D, P) and 4 types of ions (CO, K, NA, CL) are also present in the system. The complete set of interaction parameters consists of 36 non-bonded terms, 4 bonded terms and 3 angle terms. Analysis of the plots allows observation of correlations among interaction terms. For example, there would be no potential minimum for the D-P non-bonded interaction at 6.5 Å if there is no correlation with other interactions, since there is no peak at 6.5 Å in the D-P RDF.

**Figure S3.**
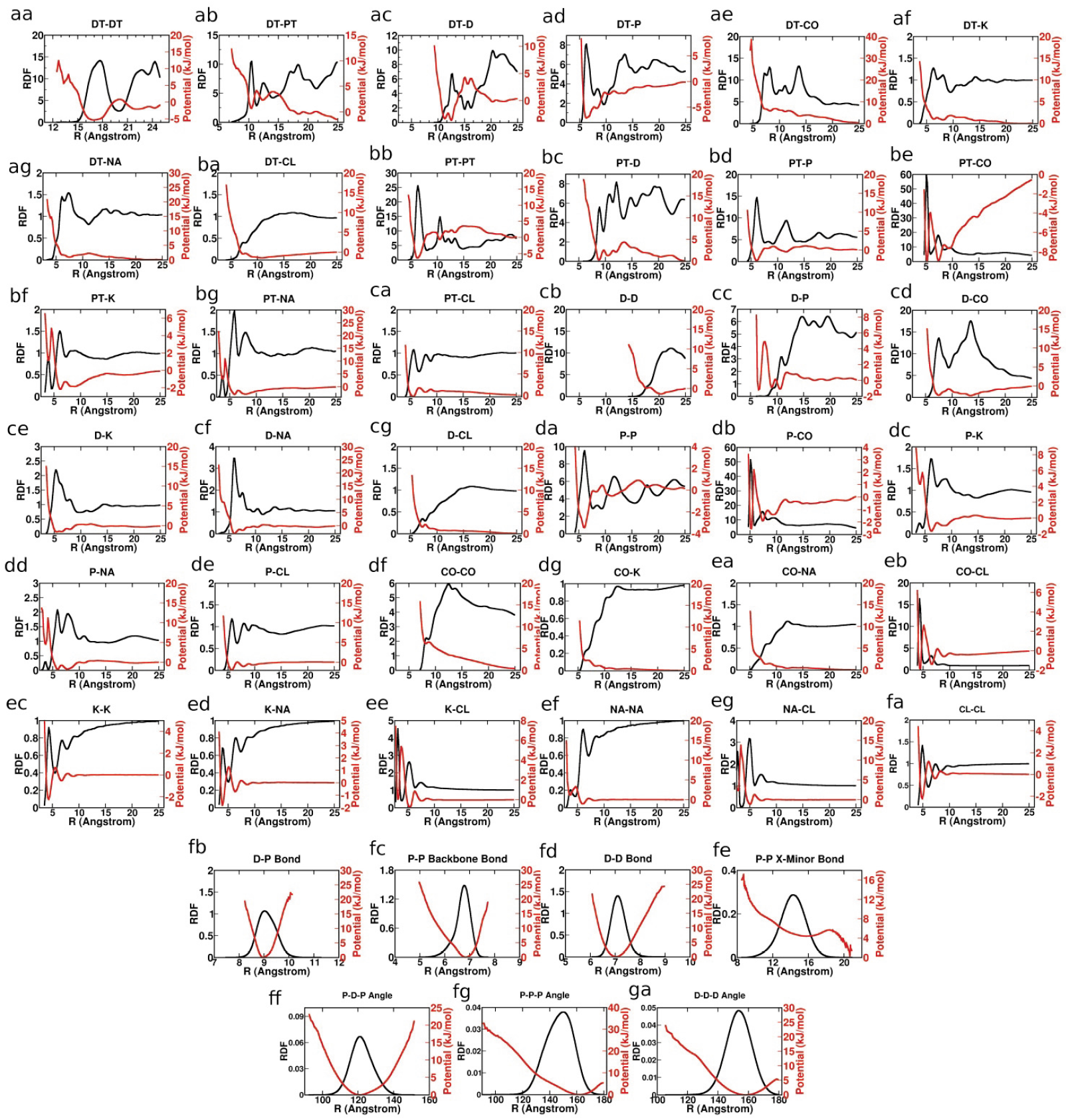
RDFs (black) and the final effective potentials (red) for all interaction terms in the CG DNA simulations with CoHex^3+^. A total of 36 non-bonded interactions (**aa – fa**), 4 bond interactions (**fb – fe**) and 3 angle interactions (**ff – ga**) are shown.

**Figure S4. Convergence of the Inverse Monte Carlo procedure.**

Examples below compare distribution functions and interaction potentials obtained in several iterations of the IMC procedure.

**Figure S4.**
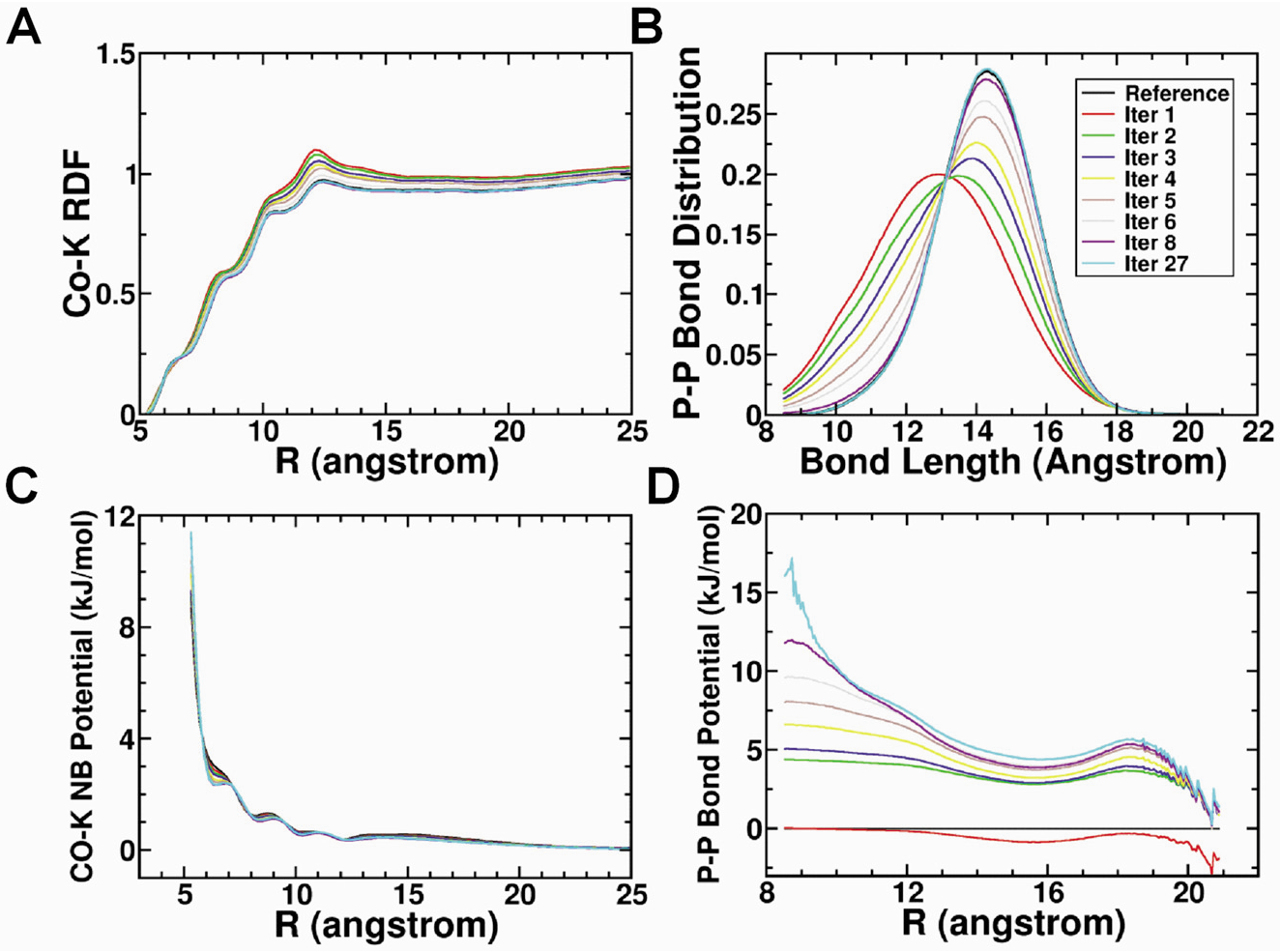
Selected RDF (**A, B**) and effective potential (**C, D**) plots from several iterations in the IMC calculation. RDFs and potentials of one non-bonded term (Co-K, plots in **A** and **C**) and one bonded term (P-P cross minor groove bond, plots in **B** and **D**) are selected to illustrate the convergence of the RDF and the effective potential. The RDF underwent significant adjustments in the first few iterations of the runs. The adjustment in the potential progressively becomes smaller and smaller. The final potential, reproducing the whole set of reference distribution functions within the computational uncertainty, is achieved after 27 iterations.

**Figure S5. Transferability of potentials from different systems I.**

Figure S5 demonstrates the transferability of similar interaction types obtained from different underlying systems with different ionic composition and macroscopic properties. It can be noted that the D-Cl potential generated from the DNA system with only NaCl ions in the all-atom setup, displays somewhat different features compared to the same ones generated from the all-atom systems containing either Mg^2+^ or CoHex^3+^. The reason is that in the NaCl-DNA system there is a limited amount of Cl^−^ in the close vicinity of DNA, especially in the groove regions (distances less than 10 Å). In case of Mg^2+^ and CoHex^3+^ counterion systems, some Cl^−^ will be brought to these locations by the presence of these multivalent counterions, which creates a "wave" in the potential at ~7.5 Å.

**Figure S5.**
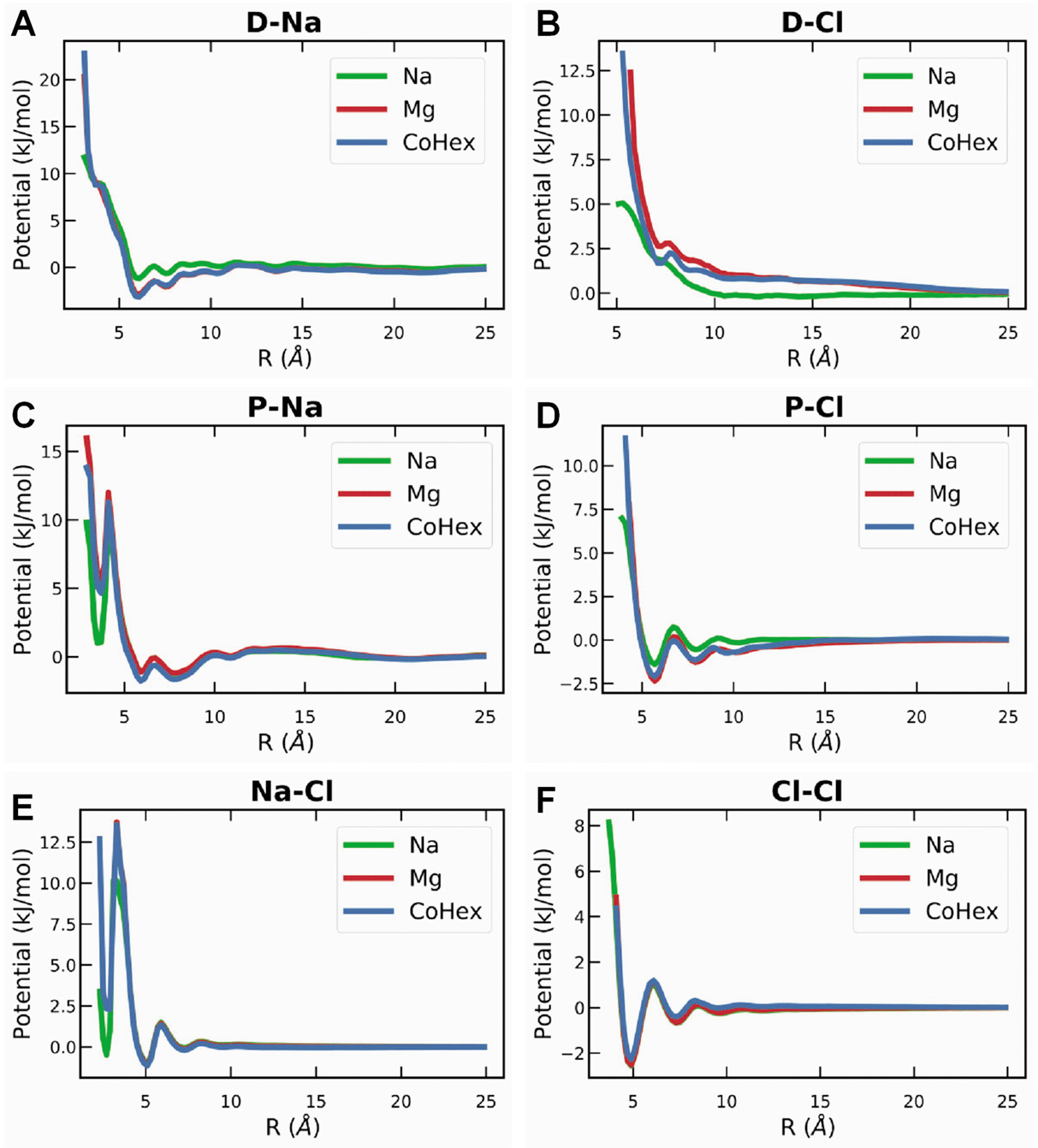
DNA-ions and ion-ion potentials obtained with the CG DNA model using the IMC procedure applied to three different DNA all-atom MD simulations of different ionic composition. (**A, B**) Potentials between DNA central bead (**D**) and respectively Na^+^ and Cl^−^ ions; (**C, D**) potentials between phosphate (P) and respectively Na^+^ and Cl^−^ ions; and (**E, F**) potentials between Na^+^ and Cl^−^ ions. Although the individual simulations display highly different behaviour, the potentials are very similar. Blue plots are obtained from the 4DNA + CoHex^3+^ simulation, green with 4DNA + Mg^2+^ simulation and the red with 1DNA + Na^+^ simulation. See Methods for details.

**Figure S6. Transferability of potentials from different systems II.**

Figure S6 below additionally demonstrates transferability of the IMC-derived potentials determined from all-atom MD simulations with different ionic conditions and number of DNA molecules in the simulation cell, resulting in very similar behaviour in macroscopic behaviour in the CG simulations. Two different all-atom systems, one with the presence of CoHex^3+^ the other Mg^2+^ were simulated (mentioned in relation to Figure 4). One displays attraction (CoHex^3+^-system) and the other displays repulsion (Mg^2+^-system). Then CG simulations for two systems containing 25 CG DNA molecules in the presence of CoHex^3+^ are conducted. One system has monovalent ion effective potentials originating from the repulsive all-atom MD Mg^2+^-system. The other system has monovalent ion potentials originating from the attractive all-atom MD CoHex^3+^-system. All other potentials (DNA and CoHex^3+^) are taken from the CoHex^3+^-system. The D–D RDF displayed in Figure S6 below illustrates that the macroscopic behaviour in these two simulations, is very similar resulting in attraction and phase separation in spite of the fact that the monovalent ion potentials are extracted from two separate all-atom MD simulations that exhibit different macroscopic behaviour.

**Figure S6.**
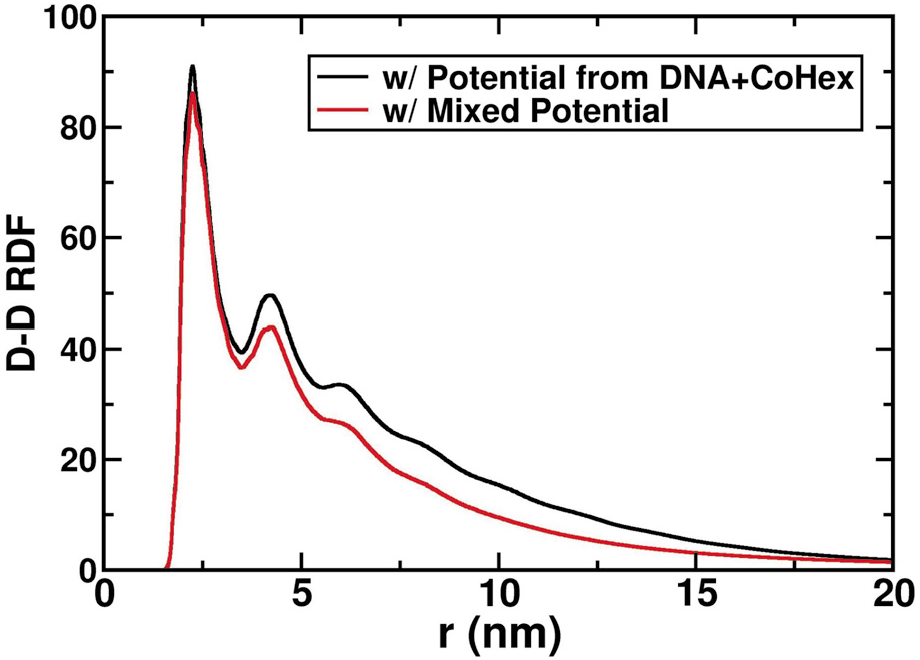
DNA-DNA distance distribution calculated for the simulations of 25 CG DNA molecules in the presence of CoHex^3+^. Black curve is the data obtained in the simulation using IMC-derived potentials from the all-atom MD simulation with CoHex^3+^ that showed DNA-DNA attraction; red curve is calculated for a system with mixed potentials where potentials for monovalent ions were taken from the Mg^2+^ - DNA all-atom MD simulation with net DNA-DNA repulsion.

**Figure S7. Statistics of toroid dimensions.**

Toroid dimensions, diameter, thickness and hole diameter, are measured in the end of each SCG simulation that have resulted in toroid formation. Example of toroid-dimension calculation is shown in Figure S8 below.

**Figure S7.**
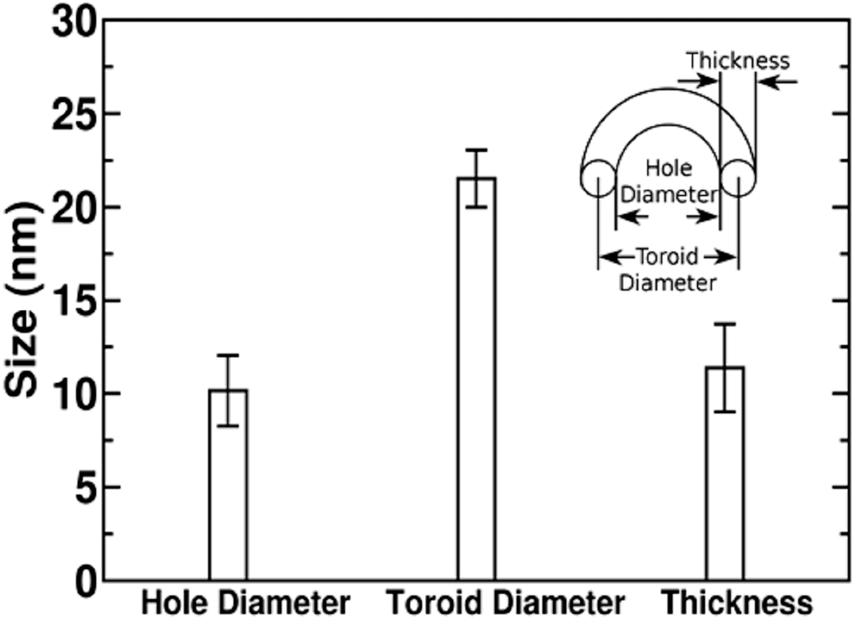
Statistics of DNA toroid dimensions. To determine the size of toroids that are formed in the simulations with the SCG model, multiple single DNA simulations have been conducted and these results are based on 17 simulations with distinct toroid formation.

**Figure S8. Measurement of toroid dimensions.**

The toroid dimensions are measured by the following method, which tries to mimic the toroid measurement practice in EM experiments. First, the toroid is projected to a plane along the axes parallel to its hole. The projected density map is used to determine two toroid dimension parameters, hole diameter and outer diameter. As shown in Figure S7B, two circles are used to represent the toroid hole and the outer dimension. The diameters of these two circles are recorded. Finally, the toroid diameter is determined to be the arithmetic average of the diameters of the two circles. Toroid thickness is defined as half the difference of the diameters of two circles.

**Figure S8.**
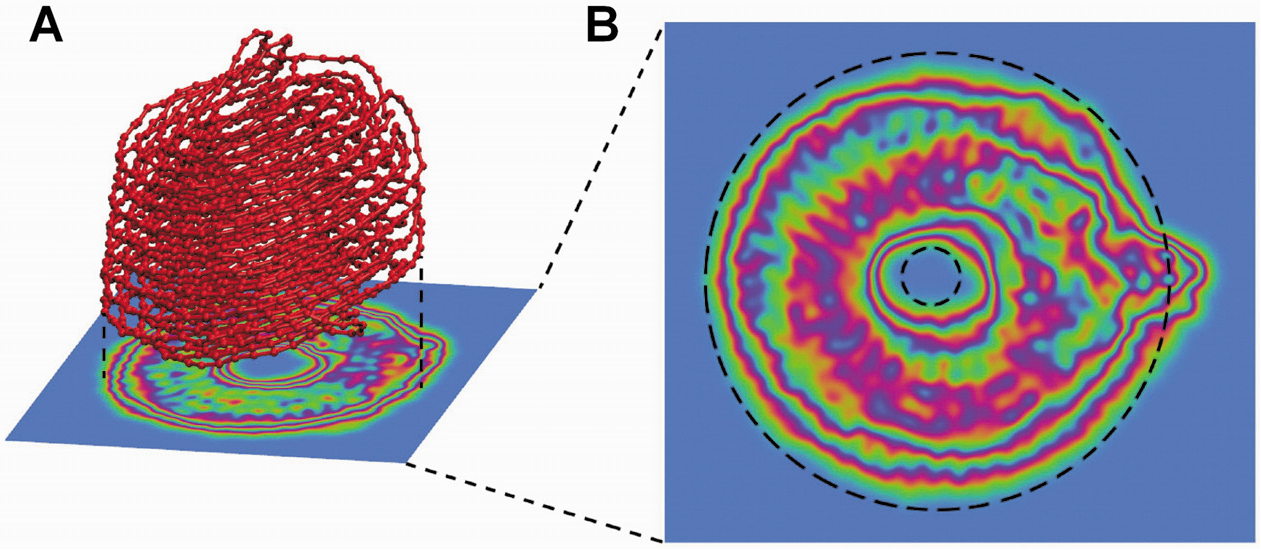
Measurement of toroid dimensions. First, the toroid is projected to a density map, on a plane perpendicular to the axes through the hole, as shown in (A). Then the projected density map was measured using two circles, one to fit the outer perimeter and one to fit the hole (B). The toroid diameter is an arithmetic average of the diameters of the two circles. Toroid thickness is the difference of the radii of the two circles.

**Supplementary Table S3**

**Table S3.** Description of intermediate states during formation of DNA toroid and fibre (see Figure 7 of the main text).

| Snapshot index (Figure 7) | Description of structure |
| --- | --- |
| **a** | Extended chain. |
| **b** | “Hairpin”: there is a turn at the end of DNA, forming a self-contact part consists of two chains of DNA. |
| **c** | Three-chain bundle: a bundle of three chains of DNA is formed at the end of DNA. |
| **d** | Thick fibre: a bundle of DNA, thicker than **c**, formed at the end of DNA. |
| **e** | “Racket”: usually on one end of the thick fibre, there’s a loop formed by at least two chains of DNA. The shape is similar to a tennis racket without string. |
| **f** | Final long and thick fibre: the final fibre we observed, usually longer than **d** and thicker than **c**. |
| **g** | Double-DNA chain loop: a loop consists of double DNA chains, usually formed from **b** when the loop in **b** folding back to the double-chain part. |
| **h** | Bundle with a side loop: the loop is formed either by newly recruited DNA or “peeling off” of one DNA chain from the bundle. |
| **i** | Primer toroid: toroid that has just been formed. It is usually very flexible. |
| **j** | Final toroid that is thick and sturdy. Grown from primer toroid by pulling in DNA chain. |
| **k** | Single loop: a loop consists of a single DNA chain part and a double DNA chain part. |
| **l** | Double loop: two adjacent single loops on DNA chain. |

## Notes

#### Summary of Updates

Changes in text and supplementary data due to revision process during journal revision

